# A computational network approach to identify predictive biomarkers and therapeutic combinations for anti-PD-1 immunotherapy in cancer

**DOI:** 10.1101/2020.04.25.055616

**Authors:** Chia-Chin Wu, Y Alan Wang, J Andrew Livingston, Jianhua Zhang, P. Andrew Futreal

## Abstract

**Background:** Despite remarkable success, only a subset of cancer patients have shown benefit from the anti-PD1 therapy. Therefore, there is a growing need to identify predictive biomarkers and therapeutic combinations for improving the clinical efficacy.

**Results:** Based upon the hypothesis that aberrations of any gene that are close to MHC class I genes in the gene network are likely to deregulate MHC I pathway and affect tumor response to anti-PD1, we developed a network approach to infer genes, pathway, and potential therapeutic target genes associated with response to PD-1/PD-L1 checkpoint immunotherapies in cancer. Our approach successfully identified genes (e.g. B2M and PTEN) and pathways (e.g. JAK/STAT and WNT) known to be associated with anti-PD1 response. Our prediction was further validated by 5 CRISPR gene sets associated with tumor resistance to cytotoxic T cells. Our results also showed that many cancer genes that act as hubs in the gene network may drive immune evasion through indirectly deregulating the MHC I pathway. The integration analysis of transcriptomic data of the 34 TCGA cancer types and our prediction reveals that MHC I-immunoregulations may be tissue-specific. The signature-based score, the MHC I association immunoscore (MIAS), calculated by integration of our prediction and TCGA melanoma transcriptomic data also showed a good correlation with patient response to anti-PD1 for 354 melanoma samples complied from 5 cohorts. In addition, most targets of the 36 compounds that have been tested in clinical trials or used for combination treatments with anti-PD1 are in the top list of our prediction (AUC=0.833). Integration of drug target data with our top prediction further identified compounds that were recently shown to enhance tumor response to anti-PD1, such as inhibitors of GSK3B, CDK, and PTK2.

**Conclusion:** Our approach is effective to identify candidate genes and pathways associated with response to anti-PD-1 therapy, and can also be employed for *in silico* screening of potential compounds to enhances the efficacy of anti-PD1 agents against cancer.

## Introduction

Breakthroughs in cancer immunotherapies have opened a new front in the war against cancer [1]. Instead of directly targeting cancer cells using specific inhibitors, immunotherapies stimulate and modulate the host’s immune system to eliminate cancer cells. Recently, immune checkpoint blockade (ICB), which enhances T-cell activity by inhibiting immunosuppressive checkpoint molecules such as cytotoxic T-lymphocyte-associated antigen 4 (CTLA-4), programmed cell death 1 (PD-1), and programmed cell death protein ligand 1 (PD-L1), has produced remarkably durable responses in some cancer patients. Despite these successes, only a subset of cancer patients benefits from these therapies, and rates of response vary widely among cancer types. Therefore, there is a growing need to understand the mechanisms underlying this de novo resistance, to select predictive biomarkers of therapeutic response, and to identify therapeutic targets that could extend the benefits of ICB [2-4].

Ongoing studies show that both cancer-cell-intrinsic and cancer-cell-extrinsic factors contribute to response to ICB [5, 6]. Initial immune activation requires the expression of neoantigens by cancer cells, which are encoded by somatic nonsynonymous mutations. Many studies have shown that nonsynonymous mutation burden is one of the most important determinants of responsiveness to ICB [7]. However, a considerable number of patients with high tumor mutation burden (TMB) have poor responses, and a subset of patients with low TMB can respond to ICB. Recently, both patient cohort studies [6-8] and genetically engineered mouse models [9] have shown that cancer cells can utilize their genetic and epigenetic aberrations to influence various aspects of the immune landscape, such as recruitment of immunosuppressive cells into the tumor microenvironment, stimulation of tumor resistance to T-cell attack, and deregulation of immune checkpoint molecule expression. In particular, alterations in or deregulation of multiple pathways in cancer cells, including *the MAPK/PTEN/PI3K, WNT/β-catenin, JAK/STAT, interferon-γ*, and antigen processing/presentation pathways, have been shown to be associated with resistance to ICB [3, 4, 10]. In addition, several cancer-cell-extrinsic factors involving tumor microenvironment, such as T-regulatory cells, myeloid-derived suppressor cells, macrophages [4], and microbes [11, 12], also affect ICB response.

Recently, various omics-based approaches have been undertaken to identify both tumor intrinsic and extrinsic factors which can serve as predictive biomarkes to ICB. First, several genomic factors, such as neoantigen and mutation burdens [13], mismatch repair deficiency [14], and somatic copy-number variation burden [15, 16], have been used to predict ICB response. Second, transcription-level data, such as PD-1/PD-L1 expression level [17], immune-cell infiltration profiling based on transcriptional signatures [18, 19], and transcriptional immunoscores [20, 21], have also been used for response prediction. Third, microbial taxonomic data obtained from analysis of 16S ribosomal RNA [11] can be also used for response prediction. However, none of these approaches considered relationships between tumour genes/pathways and ICB responses. Owing to the different genetic makeup of each tumor, the tumor immune landscape differs greatly not only between, but also within cancer types. Thus, elucidation of the underlying cancer gene/pathway-associated molecular mechanisms would facilitate the development of new avenues for personalized immune-intervention strategies. Moreover, although some associations between genetic alterations/gene expression deregualtion and patients’ responses to ICB were explored by recent next-generation sequencing studies, the small sample size of these studies limits their generalizability [8].

Several computational network approaches have been used to identify genes that are important in cancer and other diseases. Network “guilt-by-association” methods, which are based on the hypothesis that proximate genes in a molecular network are likely to be involved in the same cellular functions, have been widely used for annotating gene functions [22], identifying novel disease genes [23], and predicting therapeutic targets [24, 25]. Here, we developed a network guilt-by-association approach to identify genes and pathways associated with response to anti-PD-1/PD-L1 ICB (hereafter termed anti-PD-1 therapy). The drug target data can be mapped to the top genes in the prediction to identify compounds that could enhance tumor response to anti-PD1.

## Results

### Overview of the network-based approach

The efficacy of anti-PD-1therapy depends on the presentation of neoantigens by major histocompatibility complex (MHC) class I molecules on the surface of cancer cells for surveillance by cytotoxic CD8+ T cells [26]. Thus, we hypothesized that aberrations of any gene that are close to MHC class I genes in the gene network are likely to deregulate the MHC class I antigen processing and presentation pathway (hereafter termed the MHC I pathway) and affect tumor response to anti-PD1. Our proposed network-based approach is illustrated in Fig. 1.

**Fig. 1.**
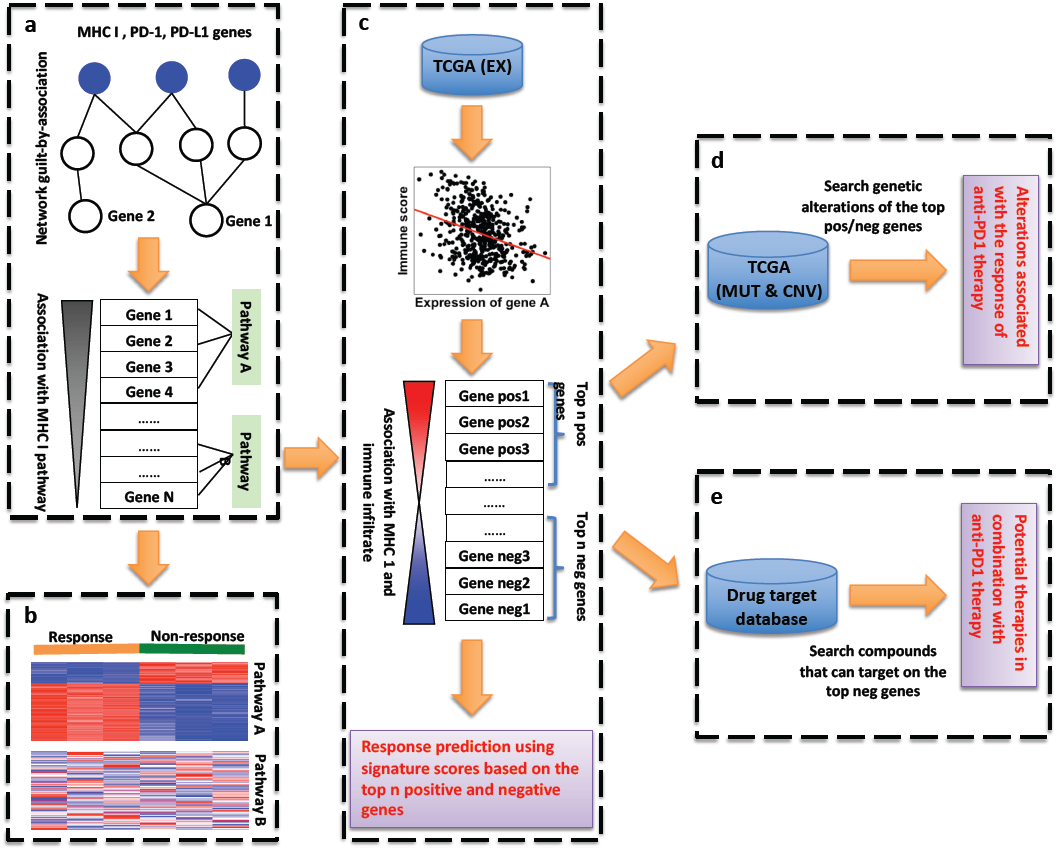
Network-based approach. **(a)** Calculation of the functional association score of each gene with MHC I genes (blue node). Genes (e.g., gene 1) that are more functionally associated with the MHC I genes are assumed to be more associated with anti-PD-1 response than other genes (e.g., gene 2) are. Gene set enrichment analysis (GSEA) was used to identify pathways associated with response to anti-PD-1 therapy on the basis of an ordered gene list ranked by the association scores. Pathway A, which enriches genes with higher association scores, is more associated with anti-PD-1 therapy than is pathway B. **(b)** Gene expression data from tumor samples that were responsive and resistant to anti-PD-1 therapy are leveraged to identify pathways associated with anti-PD-1 therapy response for a specific cancer type or cellular condition. **(c)** Integration of expression data of TCGA samples with our association predictions described in (a) to generate 2 signatures, each containing 100 top-ranked genes in our prediction list, whose expression are also mostly positively or negatively correlated with the immune infiltration score. **(d)** Identification of genomic alterations that would be associated with response to anti-PD-1 therapy in a cancer type by integrating genomic alteration data of a TCGA cancer type with the top genes described in (c). **(e)** Mapping drug target data to the top MHC I-associated genes whose expression is significantly negatively correlated with the immune score (as described in c) to identify compounds that can potentially boost response to anti-PD-1 therapy in a specific cancer type. Abbreviations: MUT, mutation; CNV, copy-number aberration.

First, the network guilt-by-association method [23, 27] was used to calculate functional relatedness (hereafter termed MHC I association score) of each genes with the MHC class I genes, PD-1, and PD-L1 (hereafter termed the MHC I pathway genes) on the basis of its relative proximity to these genes in a compiled gene network (see Methods) (Fig. 1a). Aberrations in genes that have higher the MHC I association scores are more likely to deregulate the MHC I pathway and, thus, to be associated with tumor response to anti-PD-1 therapy.

Second, we considered pathways that enrich genes with the high MHC I association scores are also associated with tumor response to anti-PD-1 therapy (Fig. 1a). Gene set enrichment analysis (GSEA) [28, 29], which can evaluate the genes in a pathway for their distribution in an ordered gene list ranked by the MHC I association scores, was used to identify pathways associated with response to anti-PD-1 therapy (termed GSEA association analysis). However, because the gene network we used in the prediction contained molecular interactions across different cellular statuses, not all of the predicted genes and pathways are associated with the MHC I pathway in a given cancer type. By leveraging gene expression data of samples from a cohort of cancer patients treated with anti-PD-1 therapy, we were able to identify genes and pathways that are both highly functionally associated with the MHC I pathway and deregulated in a given cancer type (Fig. 1b). The deregulation level of an associated pathway can be also determined using GSEA (termed GSEA deregulation analysis). The truncated product method (see Methods) was used to combine the P value of each pathway generated from the GSEA association and GSEA deregulation analyses to identify pathways that are associated with response to anti-PD-1 in a given cancer type.

Third, TCGA transcriptomic data that have been used to study the tumor immune infiltration [31] can also be integrated with our network association prediction (Fig. 1c) to identify genes and pathways that are associated with tumor response to anti-PD-1 therapy in a given cancer type. We reasoned that expression of the top genes in our prediction, which are negatively correlated with an immune infiltration score (ESTIMATE) [31], would be negatively associated with response to anti-PD-1 therapies, and vice versa. Thus, we first determined the correlation of each gene’s expression level with the immune infiltration score in a TCGA cancer type. We then integrated the resulting correlation with the MHC I-association prediction to select top immune-positive and -negative genes, which are the top 10% genes in our prediction and whose expression levels were respectively most strongly positively or negatively correlated with the immune infiltration score (absolute correlation ≥ 0.2) in a given cancer type. These genes and their pathways would be associated with tumor response to anti-PD-1 therapy in a given cancer type. A meta-analysis method by integration of our prediction and TCGA transcriptome data (see the Methods) was also used to respecitively select the top 100 immune-positive and -negative genes to calculatethe the signature score, termed the MHC I association immunoscore (MIAS), to predict patient response to anti-PD-1 therapy for a given cancer type (Fig. 1c). A patient sample with a higher MIAS score would be more likely to respond to anti-PD-1 therapy than those with a lower score. The calculation of the MIAS score is detailed in the method section.

Fourth, our MHC I-association prediction can also be integrated with TCGA genomic data (e.g., copy-number alterations and mutations) to identify genomic alterations that are associated with response to anti-PD-1 therapy in a cancer type (Fig. 1d). Here, we selected recurrently alterated genes that were in the top 10% of our prediction list and whose gene expression was also significantly positively or negatively correlated (absolute correlation ≥ 0.2) with the immune score across the samples in a cancer type.

Fifth, we also reasoned that inhibition of the top immune-negative genes in our prediction were likely to be able to boost tumor response to anti-PD-1 therapy. Therefore, drug target data compiled from the DGIdb database [32] were mapped to the top 10% of genes in our association prediction list, whose expression was also significantly negatively correlated with the immune score (correlation ≤ −0.2) in a given cancer types, to identify potential inhibitors that could enhance tumor response to anti-PD-1 therapy (Fig. 1e).

### Prediction of genes and pathways associated with MHC I pathway and response to anti-PD-1 therapy

Our approach first predicted genes that are associated with MHC I pathway genes. Some genes that are known to be directly involved in the MHC I pathway, such as *TAP1, TAP2, CALR, TAPBP*, and *B2M* [33], were at the top 1% of our prediction list (Additional file 2: Table S1). Alterations of some top genes in our prediction, such as *B2M, JAK1, JAK2, HSP90*, and *IFNG*, are known to be associated with poor response to anti-PD-1 therapy [7, 33-36]. Several genes that were recently found to be associated with anti-PD-1 response were also identified in the top list of our prediciton, such as *KRAS* [37], *STK11* [38], *DDR2* [39], and *ATR* [40].

Several groups have developed T-cell-based CRISPR/Cas9 screens to identify essential genes associated with the interaction between cancer cells and T cells [41-44]. Therefore, we used 5 gene sets from these studies to comprehensively evaluate our predictions. The receiver operating characteristic (ROC) curves and area under the curve (AUC) values for the 5 gene sets are shown in Fig. 2a. The high AUC values (> 0.7) indicated that many of the top genes in our prediction list would be highly associated with response to anti-PD-1 therapy. By pairwise overlap analysis, we also found that the top 10% of genes in our prediction significantly overlapped with all 5 CRISPR-based gene sets (Additional file 1: Fig. S1). However, the overlap among some CRISPR gene sets from the different studies was not significant. This may result from the use of different techniques or cell sources in their experiments.

**Fig. 2.**
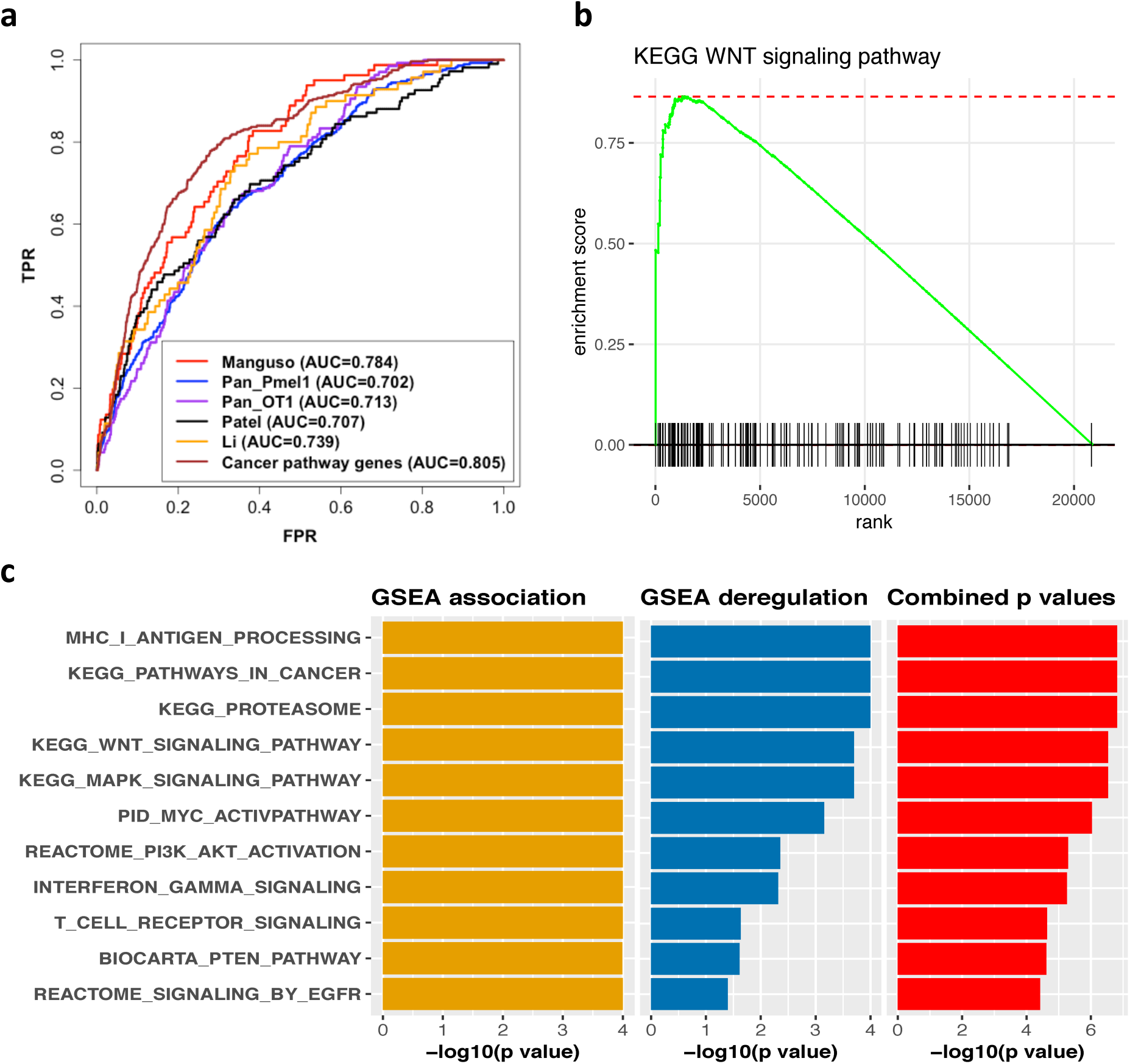
Evaluation of the MHC I-association prediction using benchmark gene sets and known pathways associated with anti-PD-1 therapy. **(a)** Receiver operating characteristic (ROC) curves for 4 CRISPR-based gene sets and a set of cancer pathway genes: 81 genes from Manguso (Manguso), 326 genes identified with Pmel-1 T cells from Pan (Pan_Pmel1), 175 genes identified with OT-1 T cells from Pan (Pan_OT1), 109 genes from Patel (Patel), and 328 genes from the Kyoto Encyclopedia of Genes and Genomes (KEGG) cancer pathways (cancer pathway genes). Genes in these 5 gene sets were treated as positive instances, and the remaining genes in the gene network were treated as negative instances. TPR indicates true positive rate; FPR, false positive rate; AUC, area under the ROC curve. **(b)** GSEA association plot of the Wnt pathway from our prediction. GSEA evaluated the genes of the pathway for their distribution in the ordered gene list generated by our association-prediction. **(c)** Some pathways that were significantly associated with the MHC I pathway in our prediction. The length of a bar represents the magnitude of the statistical significance of a given pathway in GSEA association analysis, GSEA deregulation analysis, or combination analysis using the truncated product method. The statistical significances are shown as -log10(*p*-value).

Next, we applied GSEA to the ordered list of genes in our prediction to infer pathways that are associated with response to anti-PD-1 therapy (Fig. 1b). The result also revealed that several pathways known to be associated with response to anti-PD-1 therapy were at the top of our prediction list (Fig. 2c): the WNT pathway (the GSEA association plot for this pathway is shown in Fig. 2b), *PI3K-AKT* pathway, MYC pathway, MAPK pathway, interferon signaling pathway, and TP53 pathway [45-49]. Because these association predictions were not specific to a particular cancer type, we also integrated our predictions with gene expression data of renal cell carcinoma samples from anti-PD-1 therapy responders and nonresponders [50]. We analyzed the gene expression data and also examined the deregulation of these pathways using GSEA. The combination of the GSEA association and GSEA deregulation analyses revealed pathways that are both highly associated with MHC I pathway in our prediction and significantly deregulated between response and non-response renal cell carcinoma samples (Fig. 2c).

In addition, several pathways that have been recently proposed as mechanisms of response or resistance to anti-PD-1 therapy were also identified by our approach (data not shown), such as cellular metabolism [51], DNA damage/repair pathways [52], and defective transcription elongation [53].

### Some cancer genes acting as network hubs are associated with response to anti-PD-1 therapy

We found many known cancer genes to be functionally associated with the MHC I pathway in our prediction, including *AKT2, EGFR, TP53, PTEN, MAPK1, MAPK3, CTNNB1, AKT1, JAK1*, and *JAK2* (Additional file 2: Table S1). The ROC curve evaluation of our prediction list using a set of 328 cancer-related genes compiled from the Kyoto Encyclopedia of Genes and Genomes (KEGG) cancer pathways also showed that many cancer pathway genes are in the top list of our MHC I association prediction (Fig. 2a). The overlap analysis showed that all the 5 CRISPR-based gene sets significantly overlapped with the KEGG cancer pathway gene set or with other cancer gene sets compiled from 2 databases (Fig. 3a). These results indicate that many cancer genes and their related pathways may be strongly associated with the MHC I pathway and patients response to anti-PD-1 therapy.

**Fig. 3.**
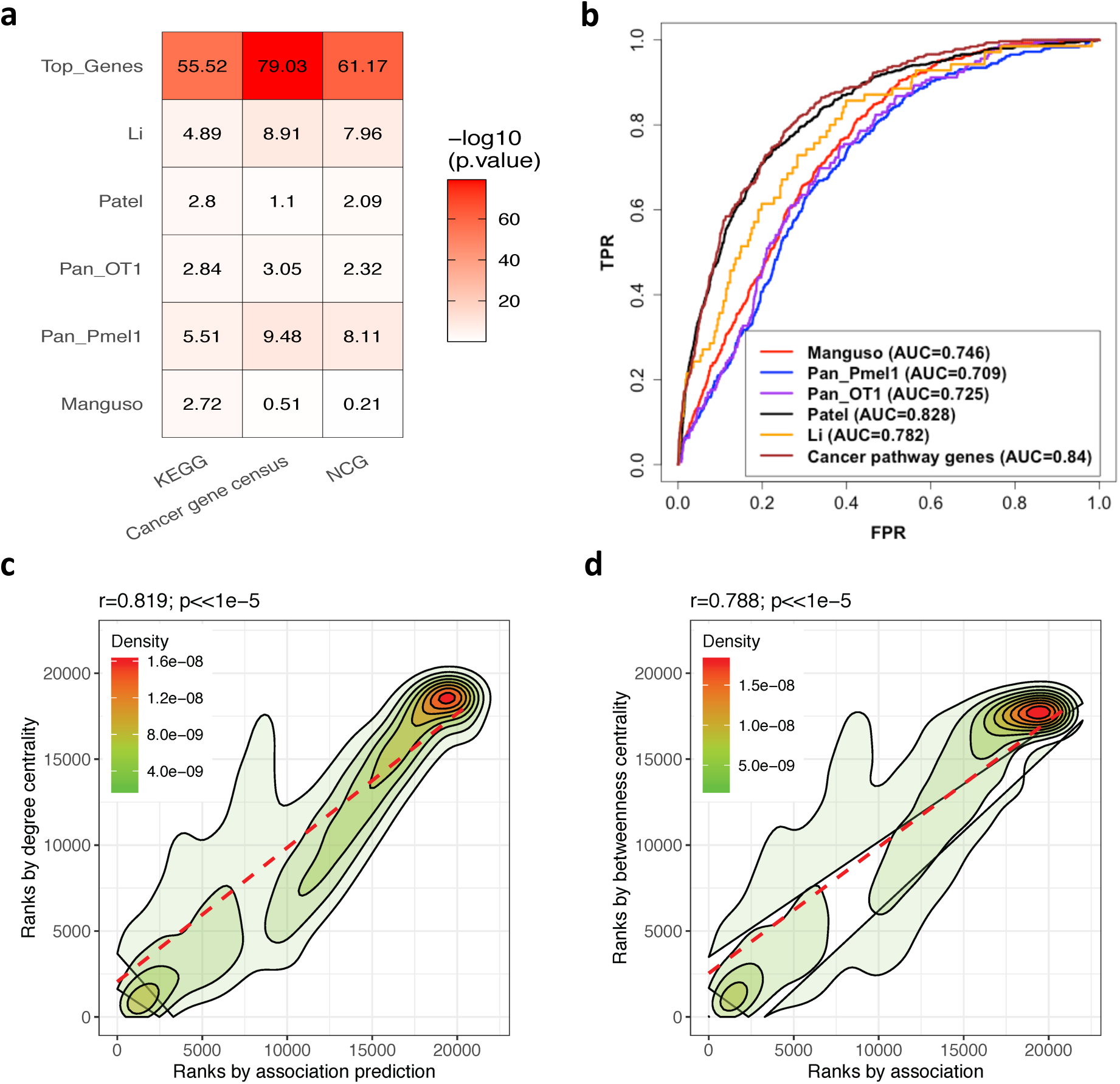
Some cancer genes acting as network hubs are associated with response to anti-PD-1 therapy. **(a)** The overlap analysis of the top 10% of genes in our MHC I-association prediction list (Top_Genes) and all the 4 CRISPR-based gene sets (same denotations as Fig. 2) with the four cancer gene sets compiled from the databases (see Methods). The color scale in the heat map graph indicates the statistical significance of the overlap, –log10(*p*-value), calculated by a hyper-geometric test. **(b)** Evaluation of predictions generated by the degree centrality method using the 4 CRISPR-based gene sets and the KEGG cancer pathway gene set. The gene sets are indicated as in Fig 2. **(c)** Correlation analysis of the prediction generated through MHC I genes and the prediction generated using the total degree centrality method. **(d)** Correlation analysis of the prediction generated through MHC I genes and the prediction generated using the betweenness centrality method.

We reasoned that these cancer genes may act like hubs in the network to directly or indirectly deregulate multiple molecular pathways [24, 54], including the MHC I pathway, to suppress immune system and promote the growth, survival, and proliferation of cancer cells. To validate this, we first calculated 3 kinds of hub centrality scores for all the genes in the compiled network: degree, betweenness, and eigenvector centrality (see the Methods). We then used the 5 CRISPR-based gene sets to evaluate the predictions generated by the 3 centrality methods. The ROC curve in Fig. 3b shows that the prediction of degree centrality correlated well with the results of all 5 CRISPR gene sets. Similar results were also observed for the other 2 centrality methods (Additional file 1: Fig. S2 and S3). Fig. 3c, 3d, and S4 (Additional file 1: Fig. S4), respectively, also show that the predictions of the degree, betweenness, and eigenvector centrality methods were all highly correlated with our MHC I association prediction. Taken together, these results suggest that some cancer genes act as hubs in molecular networks to indirectly deregulate the MHC I pathway and drive immune evasion of cancer cells.

### The MHC I-association immunoscore for predicting patient response to anti-PD-1 therapy

The integration of our MHC I-association predictions and TCGA transcriptomic data can also generate gene signatures to predict patient response to anti-PD-1 therapies in a specific cancer type (the method was detailed in the Method section) (Fig. 1c). In this work, we demonstrated this capability by applying our approach to melanoma, for which the most anti-PD-1 therapy cohorts are available. The gene expression data of TCGA skin cutaneous melanoma (SKCM) samples were first integrated with our MHC I association prediction to generate a positive and a negative gene signature (Additional file 2: Table S2). We next applied these two signatures to analyze gene expression data sets of 354 samples compiled from 5 melanoma cohorts [20,55-58], in which patients were treated with anti-PD-1 therapy alone or in combination with anti-CTLA-4 therapy, and calculated the MIAS score of each sample. The MIAS scores and the clinical response data of the samples were then used to calculate the AUC to quantify the predictive performance of our approach. This evaluation was applied to each cohort dataset individually as well as the combined dataset that merged the datasets from all cohorts. Fig. 4a shows that the AUC values of most of the data sets were substantially higher than the random expectation (AUC=0.5). However, since size of some data sets are small, the ROC curve evaluation may not be reliable [59]. Thus, we also used the Wilcoxon-Mann-Whitney statistic, which is directly connected to the AUC of a ROC curve [60], to evaluate the performance of our predictions. Indeed, we found the results of the Wilcoxon tests and the AUC to be inconsistent in some small data sets; that is, some data sets had very high AUC values but nonsignificant *P* values on the Wilcoxon test (e.g., the Auslander.PD1.On_2018 data set).

**Fig. 4.**
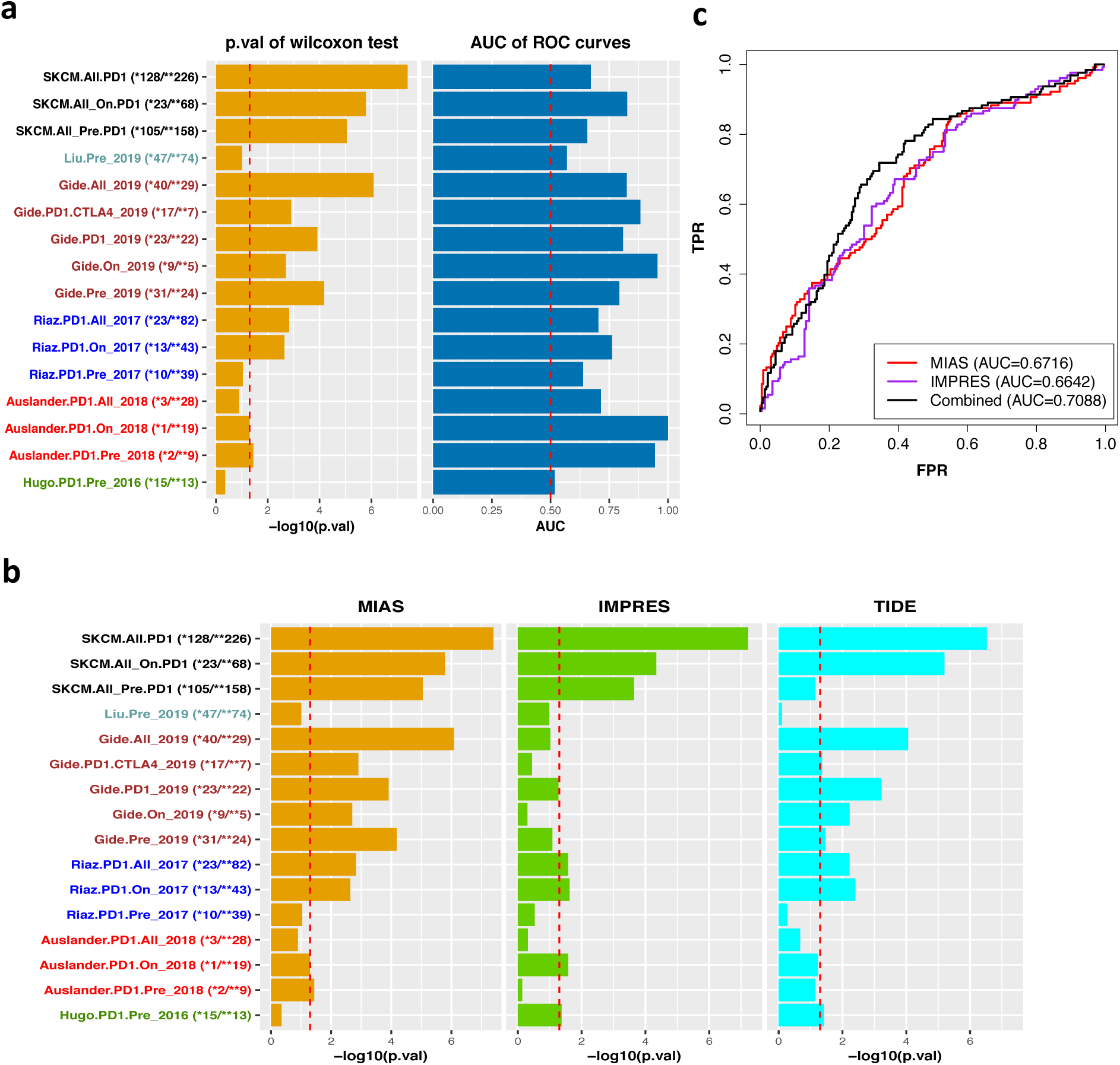
Performance of predictions of response to anti-PD-1 therapy. **(a)** *p*-values of Wilcoxon test and area under the curve (AUC) values of receiver operating characteristic (ROC) curves quantifying the predictive performance of our melanoma response predictor of anti-PD1 across several melanoma patient cohort data sets: Ascierto, Auslander, Gide, Hugo, Riaz, and the aggregated datasets (SKCM.All). ‘Pre’ indicates before treatment; ‘on’, during treatment. The vertical dotted lines in the 2 bar plots respectively represent P= 0.05 and AUC = 0.5. **(b)** Performance comparison of our approach with two other methods, IMPRES and TIDE, using Wilcoxon test across several melanoma patient cohort data sets. The dotted line in the bar plots represents *p* = 0.05. **(c)** Response prediction of the combined dataset (merged the datasets from all cohorts) was improved by integrating prediction of our approach and IMPRES.

We next compared the performance of our approach with that of 2 other recently proposed methods, IMPRES [20] and TIDE [21], by using the Wilcoxon-Mann-Whitney test (Fig. 4b) and AUC (Additional file 1: Fig. S5). The results showed that the predictive performance of our approach was a little better than those of the two other methods in most of the data sets. This may be due to the fact that our method was designed specifically based on MHC I pathway-association. In addition, all the three methods performed poor in pre-treatment datasets (AUC=0.656 for all the pre-treatment samples), except the Gide data set (AUC=0.793) [58], but performed significantly better in on-treatment datasets (AUC=0.826 for all the on-treatment samples). To investigate the difference between pre- and on-treatment samples, we compared their the ESTIAMTE immune infiltration scores and found that the immune infilitration level in pre-treatment datasets are all significantly lowere than in on-treatment datasets (Additional file 1: Fig. S6).

Furthermore, predictions made using our approach and IMPRES cover different aspects of immune response and suppression mechanisms. The IMPRES prediction is based on pairwise relations between the expression of selected inhibitory and stimulatory immune checkpoint genes. In contrast, our method considers deregulation of genes and pathways that are functionally associated with the MHC I pathway. Therefore, we reasoned that integration of the predictions of the 2 approaches may improve the overall predictive performance. We used the AUC to evaluate the prediction performance of the integrated prediction for the combined dataset. Fig. 4c shows that the performance of the integrated prediction was better than that of the 2 individual approaches. In contrast, we did not see the performance was improved in the integrated prediction of our approach and TIDE (Additional file 1: Fig. S7).

### Mining TCGA data to identify genetic aberrations associated with anti-PD-1 response

At present, the sizes of clinical cohorts treated with anti-PD-1 therapy are still very limited, making genetic association analyses of response to anti-PD-1 statistically underpowered [8]. Some large-scale cancer genome projects, such as TCGA, have revealed the genomic and transcriptomic landscapes of several cancer types. Thus, mining TCGA data using our MHC I-association prediction would allow us to explore genetic, epigenic, and transcriptomic aberrations associated with anti-PD-1 therapy in a given cancer type. By integrating the TCGA transcriptomic data, we first found that the top 10% of genes in our association prediction list significantly overlapped with those genes whose expression was significantly correlated (absolute correlation ≥ 0.2) with the immune infiltration score (ESTIMATE) [31] in most of cancer types (Fig. 5a), indicating enrichment of genes associated with immune infiltration in our top prediction list. These overlapping genes are expected to be associated (either positively or negatively) with response to anti-PD-1 therapy in each cancer type. On average, about 400 overlapping genes were identified in a cancer type; in total, 2019 genes were identified across all cancer types (Additional file 2: Table S3). Pathway analysis of the overlapping genes in each cancer type also revealed several well-known pathways associated with immunotherapy resistance in some cancer types, such as the PI3K signaling, IFN-γ, and Wnt pathways (Additional file 2: Table S4).

**Fig. 5.**
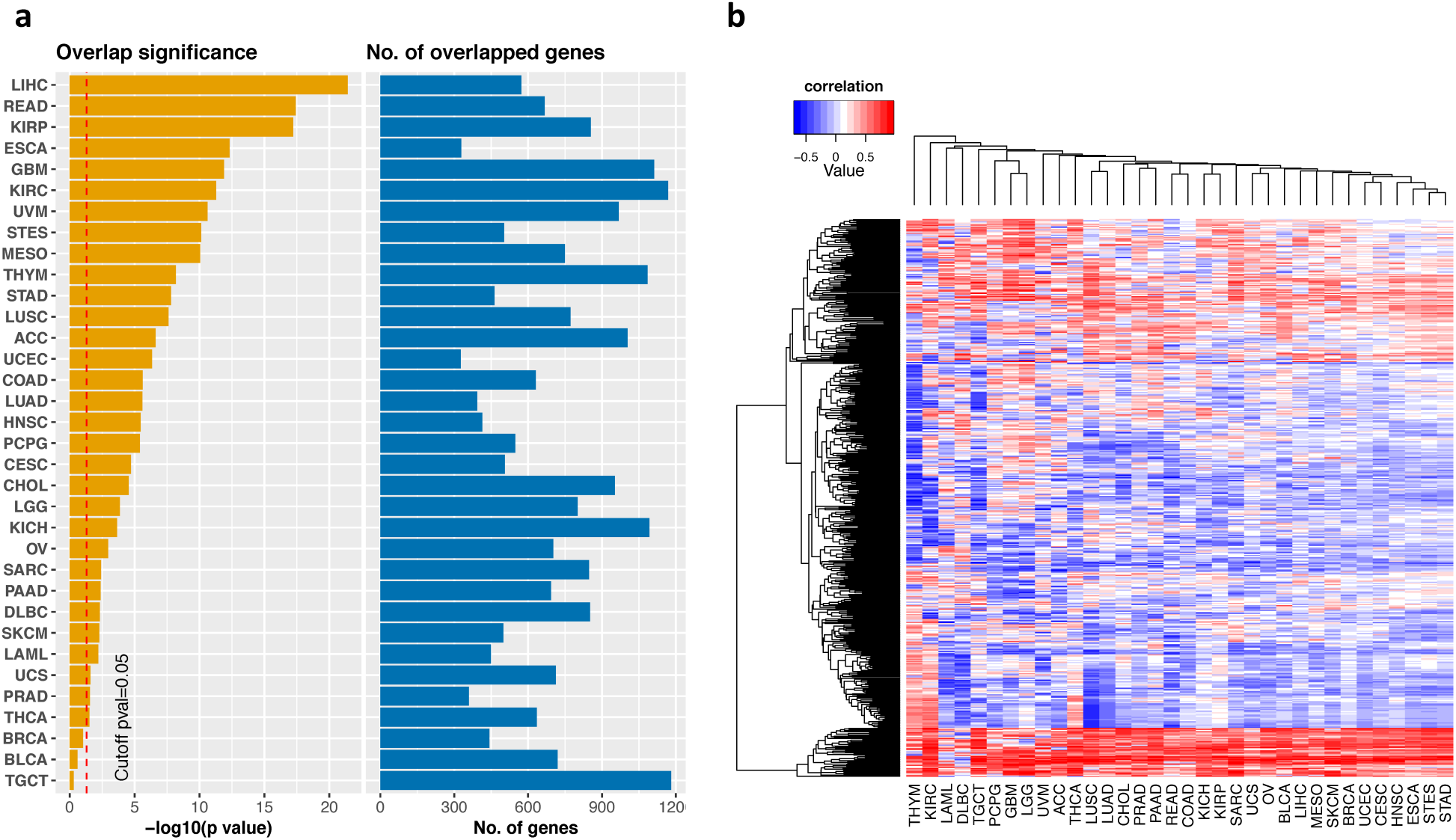
Integration of the MHC I-association prediction with TCGA data. **(a)** Overlap between the top 10% of genes in our MHC I-association prediction list and those genes whose expression is significantly correlated with immune infiltration score (absolute correlation ≥ 0.2) for the 34 TCGA cancer types. **(b)** The hierarchical clustering of cancer types using the 500 overlapping genes with the most variable immune infiltration correlation values across the 34 TCGA cancer types, revealing that cancer types in related tissue lineages were clustered together. The heatmap of all 2019 overlapping genes across all 34 TCGA cancer types (Additional file 1: Fig.S8) also reveal the similar clustering.

After comparing the overlapping genes across all cancer types, we found that only 106 of these genes were shared among all cancer types. Expression levels of all 106 genes were positively correlated with immune infiltration; most of them are related to the T-cell-induced cytolytic process. In addition, we also found that the direction of the correlation with immune infiltration for some genes was not consistent across all cancer types; that is, thier correlations were positive in some cancer types but negative in others (Fig. 5b). Furthermore, hierarchical clustering of the immune infiltration correlation of the 500 most variable genes of these 2019 genes revealed that cancer types in related tissue lineages were clustered together (Fig. 5b). For example, glioblastoma multiforme (GBM) and low-grade glioma (LGG) were clustered together, as were lung squamous cell carcinoma (LUSC)/lung adenocarcinoma (LUAD); kidney chromophobe carcinoma (KICH)/kidney renal papillary cell carcinoma (KIRP); and stomach adenocarcinoma (STAD)/stomach and esophageal carcinoma (STES). These results indicate that the molecular mechanisms associated with response or resistance to anti-PD-1 therapy may be tissue and lineage dependent.

Furthermore, we can map these overlapping genes to point mutation and copy-number data in TCGA to identify genetic aberrations that may be associated with response or resistance to anti-PD-1 therapy in each cancer type. Table 1 lists some of the identified genetic aberrations in SKCM. Several known genetic aberrations associated with resistance to anti-PD-1 therapy, such as copy-number loss of *B2M* and *IFNGR1* [35] and copy-number gain of *MYC* [46], were identified. Several identified genes were recently shown to be associated with adaptive immune response in other cancer types and would be potential biomarker and therapeutic targets in melanoma. First, The p16-cyclin D-CDK4/6-retinoblastoma protein pathway is dysregulated in most melanomas [61]. Recently, cyclin-dependent kinase 4 (*CDK4*) was found to be associated with antitumor immunity in murine models of breast carcinoma [62], and inhibition of CDK4 can increase PD-L1 expression and enhance the efficacy of anti-PD-1 therapy *in vivo* [63]. Therefore, *CDK4* copy-number gain would be expected to be associated with resistance to anti-PD-1 therapy in melanoma. Second, overexpression of protein tyrosine kinase 2 (PTK2) has been shown to promote metastasis of human melanoma xenografts [64]. Several studies also showed that silencing of *PTK2* can render pancreatic cancers responsive to checkpoint immunotherapy [65]. Copy-number gain of *PTK2* would thus be associated with resistance to anti-PD-1 therapy in melanoma. Third, ubiquitination plays important roles in the MHC I pathway [66] and in cancer progression [67]. Thus, copy-number alterations of ubiquitin C (*UBC*) may be associated with resistance to anti-PD-1 therapy in melanoma. Fourth, some studies have revealed that translation initiation factor 3b (*EIF3B*), a key subunit of the largest translation initiation factor that acts to ensure the accuracy of translation initiation, is closely related to oncogenesis [68]. Xu et al. [69] reported that *EIF3B* can accelerate the progression of esophageal squamous cell carcinoma by activating β-catenin signaling, which can promote immune escape and resistance to anti-PD-1 therapy [4]. Thus, copy-number gain of *EIF3B* may be associated with resistance to anti-PD-1 therapy in melanoma thrugh β-catenin activation.

**Table 1.**
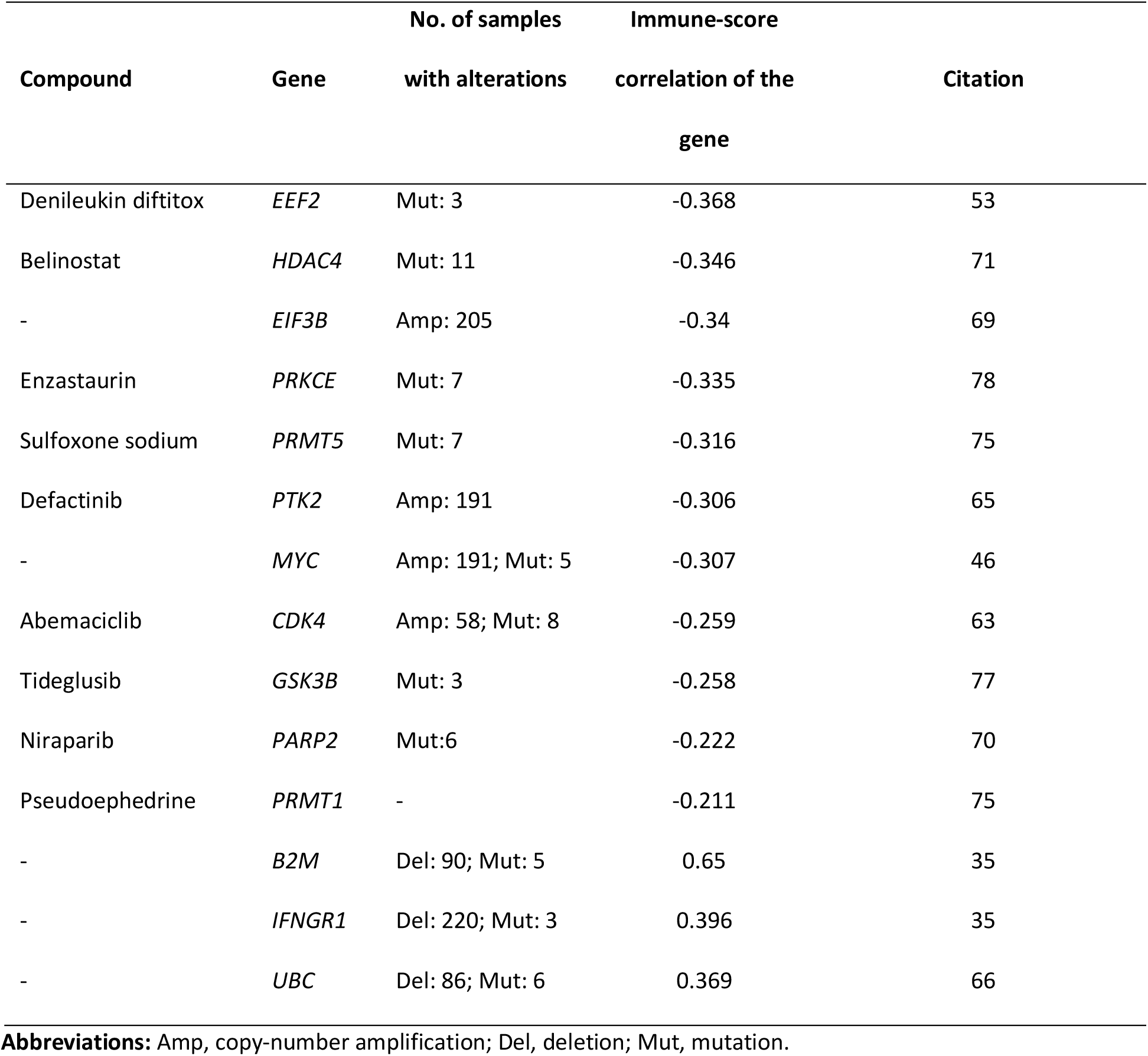
Compounds, genetic alterations, and immune-infiltration score correlations of the selected genes identified for TCGA SKCM.

### Potential therapeutic targets for combination therapy with anti-PD-1 therapy

We also reasoned that therapies that target some of the top genes and pathways in our prediction list may overcome resistance and broaden the clinical utility of anti-PD1 therapy. Recently, several clinical trials of targeted therapies and chemotherapies in combinations with anti-PD-1 therapies have been performed. To evaluate our method’s ability to predict therapeutic targets for uses in combination with anti-PD-1 therapies, we manually collected 155 target genes of 36 compounds that that have been tested in clinical trials or used for combination treatments with anti-PD-1 therapies in cancer. Fig.6 shows that many of the targets of these compounds are in the top prediciton lists of our MHC I-association analysis and the 3 centrality methods. This indicates that these network approaches may help identify targeted therapeutic strategies to enhance patient response to anti-PD-1 therapies.

**Fig. 6.**
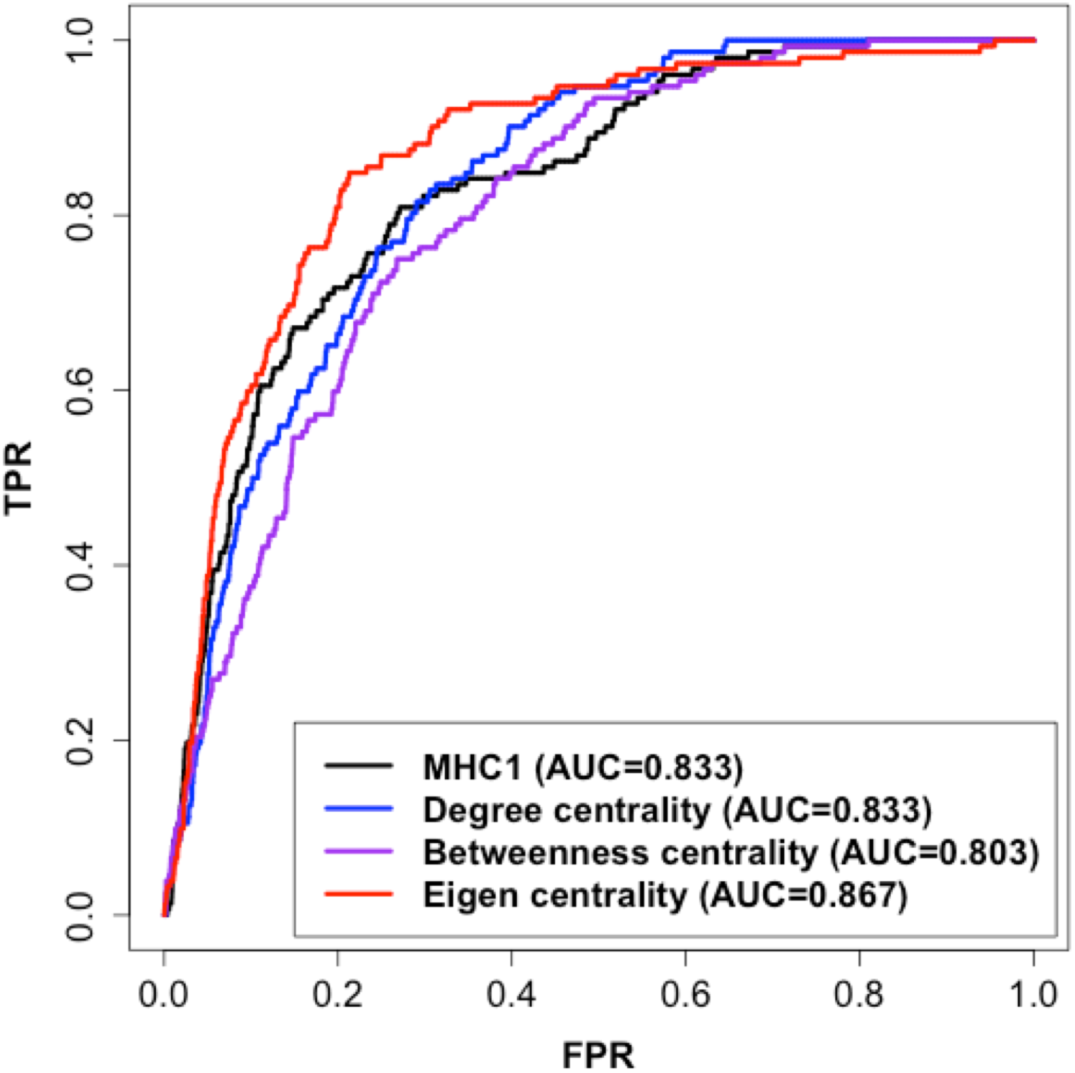
Prediction performance of therapeutic targets for combination treatments with anti-PD-1. ROC curve evaluating the MHC I-association approach using target genes of 36 compounds that have been tested in clinical trials or used for combination treatments with anti-PD-1 therapy, indicating that our network approach can help identify targeted therapeutic strategies to enhance patient response to anti-PD-1 therapies.

We also explored TCGA data sets to identify additional potential therapeutic targets in TCGA SKCM. We selected the top 10% of genes in our MHC I-association prediction list whose expression was also significantly negatively correlated with immune infiltration in SKCM. Inhibition of the expression of these genes would be hypothesized to enhance immune infiltration, making them potential therapeutic targets for combination treatments with anti-PD-1 therapy. Drug target data compiled from the DGIdb database [32] were then mapped to these selected genes to identify potential compounds for use in combination treatment with anti-PD-1 therapy. Table 1 shows some of the identified compounds for SKCM and citations to the literature associating them with immune response. Some of our identified compounds, such as histone deacetylase (HDAC) and poly ADP ribose polymerase (PARP) inhibitors, which were not included in the 36 compounds used for the performance evaluation, are now in ongoing clinical trials in combination with anti-PD-1 therapy [70, 71]. Other identified compounds whose targets have been shown to be associated with immune response could also be used in combination treatment with anti-PD-1 therapy. First, denileukin diftitox, an antineoplastic agent used to treat leukemia and lymphoma, inhibits protein synthesis by ADP ribosylation of elongation factor 2 (EEF2), resulting in cell death [72]. A recent study [53] reported that overexpressed EEF2 suppresses proinflammatory response pathways and correlates with poor response in patients with renal cell carcinoma and metastatic melanoma treated with anti-PD-1 therapy. This implies that denileukin diftitox could be used to enhance immune response in melanoma. Second, protein arginine methyltransferase (PRMT1) is involved in interferon signaling [73], and a few studies have noted that protein arginine methyltransferase (PRMT5) may be involved in modulating expression of some immune response genes [74]. Several studies recently showed that some epigenetic-targeted drugs, such as DNA methyltransferase inhibitors, can induce T-cell attraction and enhance immune checkpoint inhibitor efficacy in some mouse models [75]. Thus, inhibitors of PRMT1 and PMRT5 could be used in combination treatments with anti-PD-1 therapy in melanoma. Third, Li et al. [76] showed that glycogen synthase kinase 3β (GSK3β) interacts with PD-L1 and can induce phosphorylation-dependent proteasome degradation of PD-L1. A study by Taylor and colleagues [77] also reported that GSK3β inhibitors can downregulate PD-1 expression and enhance CD8+ T-cell function in cancer therapy. Thus, the combination of tideglusib, a GSK3β inhibitor, with anti-PD-1 therapy could be a promising strategy for melanoma treatment. Fourth, protein kinase C epsilon (PRKCE) is overexpressed in most solid tumors and plays critical roles in different cancer-associated pathways, including toll-like receptor 4 (TLR-4) signaling that plays a role in the induction of both innate and adaptive immunity [78]. TLR-4 signaling was recently shown to improve anti-PD-1 therapy during chronic viral infection [79]. Enzastaurin, a protein kinase C inhibitor, thus could be used to enhance anti-PD-1 therapy in melanoma.

## Discussion

### Comparison of our association prediction and CRISPR/Cas9 and RNA interference screens

The results in this work demonstrated that our approache can effectively identify genes and pathways associatd with response to anti-PD-1 therapy in cancer. CRISPR/Cas9 and RNA interference screens recently have been applied to identify genes associated with response to immunotherapy without using any prior biological information. However, when no prior biological information is used, CRISPR/Cas9 and RNA interference screens must be used to assess all genes. In contrast, by integrating our prediction, we only need to screen the top predicted genes. Our analysis also showed that the limited overlaps among results of different CRISPR-based screens may be associated with different techniques or cell sources used. In contrast, the top genes in our prediction significantly overlapped with all the CRISPR-based gene sets, suggesting that our approach may be relatively unbiased than the CRISPR-based approaches.

### Cancer genes associated with response to anti-PD1 therapy

Our centrality analysis revealed that some cancer genes act like hubs in their molecular networks and are able to drive immune evasion through directly and indirectly deregulating the MHC I pathways. Therefore, some network centrality methods also may be used to identify genes and pathways associated with response and resistance to anti-PD1. The results of an *in vivo* study by Lesterhuis et al. [80] support this view. Lesterhuis and colleagues used analysis of gene networks inferred from gene expression data of responding and nonresponding tumors in murine models to identify hub genes and modules associated with response to anti-CTLA-4. They showed that targeting some of these identified hub genes with selected drugs dramatically enhanced the efficacy of CTLA-4 blockade in their murine models. However, the targets identified in mice may not be applicable in human tumors, and co-expression networks constructed from small samples may contain many false positives and negatives [81]. Nonetheless, this evidence shows that mutations or deregulation of some cancer-associated pathways can contribute to resistance to immunotherapies; importantly, many of these pathways are targetable.

### Combination therapies

Targeted therapies are directed against important dysregulated molecular pathways or mutant proteins that are required for the growth and survival of malignant cells. There has been a growing interest in combining target therapies with ICB due to the immune modulating effects of many targeted agents. First, targeted therapies can cause tumor death, which leads to the release of neoantigens and thus enhances the efficacy of checkpoint inhibition [82]. Second, targeted therapies can restore deregulated pathways associated with immune-suppressive mechanisms, such as T-cell dysfunction and exclusion, to overcome resistance and broaden the clinical utility of immunotherapy. In a recent study addressing the first point, Colli et al. [82] estimated the proportion of solid tumors that might benefit from immuno-targeted combination therapy. They surveyed thousands of TCGA genomic profiles for cases with specific mutations targeted by current agents and with a burden of nonsynonymous mutations that exceeded a proposed threshold for response to ICB. Unlike the work of Colli et al., our approach addresses the second point. Our prediction method also can be easily integrated with patient mutation-burden data to identify potential therapeutic targets and compounds that comply with both of the 2 aforementioned clinical benefits for immuno-targeted combination therapies.

Recently, several efforts tested possible drug combinations with checkpoint inhibitors. However, tests of all the possible drug combinations with PD-1/PD-L1 inhibitors may exceed the number of eligible cancer patients who can be enrolled in clinical trials, and are therefore not feasible in the context of clinical trials [39]. With the incorporation of drug target information, our approach can be used to effectively identify compounds that may have action in pathways that enhance response to anti-PD-1 therapy. However, precisely predicting the possible therapeutic effects of the identified drugs in the combination treatments is difficult for several reasons, including target-dependent drug-binding affinities, unclear mechanisms of drug action, and different types of phenotypic effects (positive or negative) induced by different targets of a drug [24, 83]. However, our approach could serve as a quick way to initially screen a small number of potential drugs for further evaluation and to help reduce the search space in screenings [84]. Moreover, our approach may help to reduce drug discovery cost by finding new uses of some approved drugs in the combination treatment with anti-PD1 for cancer patients (i.e., drug repurposing).

Our integrative analysis of TCGA data and our network association prediction also indicated that the molecular mechanisms associated with patient response to anti-PD-1 therapy are context dependent and tissue specific. Other studies also showed that immunoregulations are tissue specific [85]. Therefore, integrating other high-throughput data, such as gene expression data, with the top genes in our prediction list would help prioritize important biomarkers and potential therapeutic targets for combination threatments with anti-PD-1 therapy for a given cancer type. Furthermore, integrating our predictions with individual patient data (genomic, transcriptomic, or epigenetic) may help to develop personalized immuno-oncology treatments.

### Prediction of patient response to anti-PD1

We also demonstrated that integrating TCGA data with our network predictions can calculate the signature score, MIAS, which correlates with response to anti-PD-1 therapy in a given cancer type. Factors in addition to the deregulation of the MHC I pathway, which have been shown to be associated with response/resistance to anti-PD1, such as mutation burden, the microbiome, environmental factors, germline genetics, and immune infiltration, also need to be included in the response prediciton [86, 87]. Thus, development of integrative computational approaches on the basis of different cancer cell-intrinsic and -extrinsic factors would perform better than approaches that only consider an individual factor. For instance, the combination of mutation burden and tumor aneuploidy scores was shown to better predict response to checkpoint immunotherapies than either score alone [15, 16]. Our anaylsis also showed that the integration of the MIAS and IMPRES scores also have a better prediction performance than the two individual method.

In addition, our results also showed that MIAS, IMRPES, and TIDE all performed significantly worse in pre-treatment cohorts than in on-treatment cohorts, and the estimated immune infilitration levels in pre-treatment datasets are also significantly lower than that in on-treatment datasets. This suggests that anti-PD1 therapy may induce dramatic transcriptome changes in tumor microenvironment and thus increase prediction power of the three methods in on-treatment samples. In fact, the gene signatures of IMRPES and TIDES as well as our immune-positive signatures are related to the immune response and T cell activation. Similarly, Chen et al. [88] found that gene expression profile between responder and nonresponder are not significantly different for pre-treatment samples, but much more significantly for on-treatment samples. The gene expression of antigen presentation, T-cell activation, and T-cell homing are significantly upregulated in responders of those on-treatment samples. They concluded that gene expression profile in early on-treatment samples is highly predictive of response to anti-PD1 therapy, but is not robust in those pre-treatment samples. Therefore, we suggest that other types of data, such as mutation burden, may need to be integrated for more robust response prediction in pre-treatment samples.

Our analysis also showed that a large patient data sets is essential to build and evaluate a model approach robustly. However, it is still challenging to build a comprehensive prediction model by integrating various kinds of factors, mainly owing to a lack of large-scale patient cohorts with multilevel factor data [86]. Furthermore, our response predictor was built based on analysis of bulk gene expression data that mainly reflect the averaged gene expression across different types of cells, including malignant cells, normal host cells, stroma cells, a wide variety of immune cells, and other cells. Single cell RNA-seq analysis enables identification and quantification of cell populations presented in a given dataset. Therefore, integrating single cell RNA-seq data and our association network approach may be able to analyze gene expression of cancer cells and tumor cells sperately and develop more robust response predictors.

## Conclusion

While our approach cannot be used to identify biomarker genes whose molecular interactions have not yet been well characterized because it is based on incomplete molecular interaction data, it will improve over time as more data are generated [25]. Regardless, these results suggest that our approach is an effective method to identify genes/pathways associated with response/resistance to anti-PD1 therapy and can be employed as in-silico screening of potential drugs for combination regimens with anti-PD1 therapy in cancer.

## Methods

### Network guilt-by-association using random walk with restart

A gene network that was compiled from several curated databases [89-96], consisting of 20,909 genes and over 645,406 molecular interactions, was used in the network guilt-by-association analysis. Given a set of MHC class I genes as bait genes, we used the random walk with restart to calculate association scores for each gene with te MHC I pathway genes. In a network with n nodes (i.e., n genes in the gene network), the random walk with restart is defined as [27]

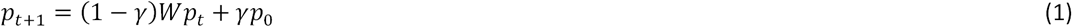

where *p*_0_ is the initial probability vector in which equal probabilities are assigned to the starting nodes (i.e., MHC class I genes, PD-1, and PD-L1); *p*_*t*_ is the probability vector containing the probabilities of the nodes at step *t*; *γ* is the restarting probability; and *W* is the transition matrix, which is a column-normalized adjacency matrix of the network. Starting from the set of nodes in the network, the walker will iteratively move from the current nodes to randomly selected neighbor nodes or return to the starting nodes. When iteratively reaching stability (i.e., when the change between *p*_*t*_ and *p*_*t*+1_ is below 10-30), the probability vector can present the association scores of all genes in the network with the starting genes. Thus, genes with higher association scores are more functionally associated with the MHC I pathway.

### Centrality measurements to identify hub genes

Network centrality measures have been used to determine hub genes in a network. These measurements also have also been used to identify genes associated with cancer and other diseases [23, 24]. In this work, 3 centrality measures—total degree [97], betweenness [98], and eigenvector centrality [99]—were used to identify genes associated with anti-PD-1 response.

In a binary network (e.g., the curated gene network mentioned above) with *n* nodes (i.e., a node represents a gene in the curated gene network), the total degree centrality of a node *i (d*_*i*_) is defined as the total number of linkages connecting it with other nodes in the network:

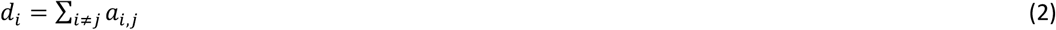

where *d*_*i*_ is the centrality measurement of gene *i*, and *a*_*i,j*_ indicates a connection between gene *i* and gene *j* in the binary network (i.e., *a*_*i,j*_ = 1 if there is a linkage between nodes *i* and *j* and *a*_*i,j*_ = 0 otherwise). Total degree centrality is the simplest way to define the centrality of a node in a network, but it only counts the local impact of a node through its direct connections in a network. Thus, some bottleneck hubs [100], which have few connections with other nodes but act as key connectors in a network, cannot be identified using total degree centrality. Thus, we also used betweenness centrality [98] in the analysis, since it can count the global importance of a node in a network through the node’s direct and indirect connections. Betweenness centrality estimates the centrality of a node in a network by considering the fraction of shortest paths that pass through that node. The betweenness centrality of a node *i* (*b*_*i*_) is defined as:

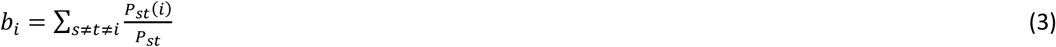

where *P*_*st*_ is total number of shortest paths between nodes *s* and *t*, and *P*_*st*_*(i)* is the number of those shortest paths between *s* and *t* that pass through the node *i*.

Another global centrality measurement used in this work is eigenvector centrality, which estimates the centrality of a node in a network based on the concept that nodes are important if they are connected to nodes that are themselves central within the network. The eigenvector centrality score of a node i (*e*_*i*_) is defined as the sum of the centrality values of the nodes that it is connected to [99]:

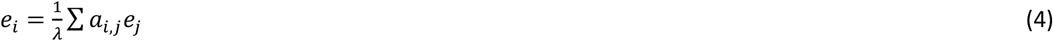

The equation can be rewritten in vector notation as the eigenvector equation

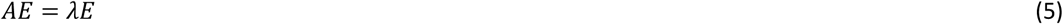

where *A* = (*a*_*i,j*_) is the adjacency matrix of the network (i.e., *a*_*i,j*_ = 1 if node *i* is connected to node *j*, and *a*_*i,j*_ = 0 otherwise), *E* = (*e*_*i*_) is a positive eigenvector, and *λ* is an eigenvalue constant.

### Identifying pathways associated with the anti-PD-1 response in a given cancer type

The truncated product method [30] was used to combine the *P* values of each pathway generated from the GSEA association and GSEA deregulation analyses to identify pathways that were both highly associated with the MHC I pathway in our prediciton and deregulated in a specific cellular condition. In the truncated product method, the product score W of the 2 *P* values (p_i_) that do not exceed a fixed τ value (τ was set to 0.01 for both GSEA association and deregulation analysis) can be calculated as:

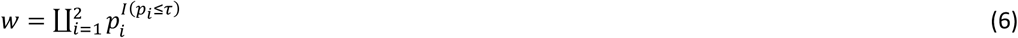

where I(.) is the indicator function. The probability of W for w < 1 can be evaluated by conditioning on k, the number of pi values less than τ:

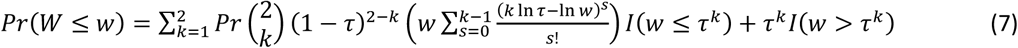

### Gene sets for performance evaluation of the network association prediction

Two types of gene sets were used to evaluate our predictions. First, T-cell-based CRISPR/Cas9 screens were recently applied to identify genes associated with mechanisms of tumor-cell resistance to killing by cytotoxic T cells. Five gene sets compiled from 4 such studies were used for evaluating our prediction, including 81 genes from Manguso’s studies (the P values in the studies <=0.05)[41], 109 genes from Patel’s studies (the P values indeitified by the RIGER metric in the studies <=0.0005) [42], 326 genes identified with Pmel-1 T cells and 175 genes identified with OT-I T cells from Pan’s studies (the P values in the studies <=0.05) [43], and 74 genes from Li’s studies (the P values in the studies <=0.05) [44]. Second, several sets of cancer-associated genes were used to evaluate the association between cancer pathways and our predictions: 328 cancer-related genes compiled from KEGG cancer pathways, 723 genes from the Cancer Gene Census [101], and 2,372 genes from The Network of Cancer Genes (NCG) [102]. Fourth, to evaluate our prediction of therapeutic targets associated with anti-PD-1 therapies, we manually collected 155 target genes of 36 compounds that have been in clinical trials and already used for immuno (anti-PD-1)-targeted combination therapies in cancer treatment (Additional file 2: Table S5). The 36 compounds were collected from several review papers [2, 103-105]. The known target genes of these compounds were compiled from the DGIdb database [32].

### TCGA data

Gene expression data for 34 TCGA cancer types were downloaded from the Broad GDAC Firehose (https://gdac.broadinstitute.org/) and were standardized across samples by quantile normalization. The immune infiltration score of each TCGA sample was calculated using ESTIMATE [31], and the correlation of each gene’s expression level with the immune infiltration score across samples of a TCGA cancer type was then calculated. Mutation calls and copy number segements of the 34 TCGA cancer types were also downloaded from the GDAC. The R package, CNTools [106] was used to convert the segment data into copy number of genes. For assessing genes with copy number aberrations, log2 values of copy number > 0.5 were considered gains while log2 values < −0.5 were considered losses.

### Prediction of anti-PD-1 therapy response and the performance evaluation

To generate the positive and negative signatures of the anti-PD-1 response for a cancer type, we first classified each gene into immune-positive or immune-negative categories on the basis of the correlation of its expression level with the ESTIMATE immune infiltration score [31] across all samples in a TCGA cancer type. For each gene *g* in each immune-correlation category *i* (positive or negtive), we used the rank product statstics [107] to merge its rank in the MHC I-association prediction (*r1*_*g,i*_) and its rank based on the positive or negative immune infiltrate correlation (*r2*_*g,i*_) to have a merged rank (*MR*_*i*_):

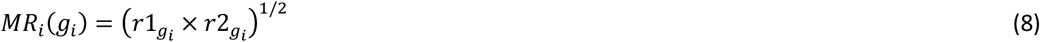

Thus, genes with the top merged ranks, *MR*_*i*_, are the top genes in our MHC I-association prediction and whose expression levels were also most strongly positively or negatively correlated with the immune infiltration score in a given cancer type. We respecitively selected the top 100 immune-positive and -negative genes based on the merged rank *MR*_*i*_ (Fig. 1c) for a cancer type. We reasoned that higher the expression levels of the top 100 immune-positive than those of the top 100 immune-negative genes are in a patient sample, the more immune infilitration the sample may have, and the more likely it can reponse to anti-PD1 therapy. Therefore, we compare the expression levels of the top 100 immune-positive genes with those of the top 100 immune-negative genes using one-tailed Wilcoxon signed rank tests for each patient sample. The MHC I association immunoscore (MIAS) of a patient sample *s* is then defined as:

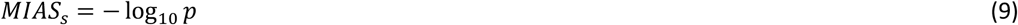

where p is the P-value of the one-tailed Wilcoxon signed rank test. A patient sample with a higher MIAS score would be more likely to have a response to anti-PD-1 therapy than would one with a lower score.

In this work, we applied this method to 354 pre- or on-treatment tumor samples (anti-PD-1 or combination of anti-PD-1 and anti-CTLA-4), compiled from 5 melanoma cohorts [20, 55-58]. The top 100 immune-positive and -negative genes (Table S2) were first selected by the integrative analysis of TCGA SKCM transcriptomic data and our MHC I-association prediction. The raw count of the RNA-seq data of the 354 smaples were convereted to transcripts per million (TPM) that normalize counts for library size and gene length. To compare the MIAS score across the samples in different cohorts, we selected the 14,820 common genes covered by all the platforms (including TCGA platform) for the MIAS score calculation. According to the clinical RECIST response information provided in the 5 cohort studies, a samples was categorized as response (complete response and partial response) and non-response (stable disease and progressive disease). Table S6 (Additional file 2: Table S6) summarizes the response annotations of the samples. Finally, the prediction performance of the MIAS score was performed using the Wilcoxon signed rank test and AUC of the ROC curve.

We also compared the performance of our method with that of 2 other recently proposed methods, IMPRES [20] and TIDE [21], using the data of the 354 melanoma samples. The IMPRES score of a sample is calculated by counting the number of the identified 15 pairwise gene expression relationships that are fulfilled. The TIDE score of a samples was calculated using a published online tool (http://tide.dfci.harvard.edu/), and the TPM values used for the calculation were further normalized as suggested in the paper. The rank product statstics [107] was used to integrate the predictions of our approach and MIAS or TIDE.

## Availability of data and materials

Other data and materials that do not appear in the main text are listed in the additional file 1 and 2 (supplementary figures and tables).

## Acknowledgement

This work was supported by Amschwand Sarcoma Cancer Foundation Award 2017 (CCW and PAF), Cancer Prevention Research Institute of Texas R120501 (PAF) and Welch Foundation’s Robert A. Welch Distinguished University Chair Award G-0040 (PAF). This work was also in part supported by NIH 1RO1CA231349-01-A1 (YAW) and Emerson Collective (YAW).

## Supplementary Figure Legends

**Fig. S1**: Heat map showing the statistical significance of the pairwise overlap analysis of the top 10% of genes in our MHC I-association prediction list (Top_Genes) and each of the 4 CRISPR-based gene sets (same denotations as Fig. 2). The color scale in the heat map graph indicates the statistical significance of the overlap, –log10(p-value), calculated by a hyper-geometric test.

**Fig. S2**: Evaluation of predictions generated by the betweenness centrality method using the 4 CRISPR-based gene sets and the KEGG cancer pathway gene set. The gene sets are indicated as in Fig 2.

**Fig. S3**: Evaluation of predictions generated by the eigenvector centrality method using the 4 CRISPR-based gene sets and the KEGG cancer pathway gene set. The gene sets are indicated as in Fig 2.

**Fig. S4**: Correlation analysis of the prediction generated through MHC I genes and the prediction generated using the eigenvector centrality method.

**Fig. S5**: Performance comparison of our approach with two other methods, IMPRES and TIDE, using area under the curve (AUC) values of receiver operating characteristic (ROC) curves across several melanoma patient cohort data sets. The dotted line in the bar plots represents AUC = 0.5.

**Fig. S6**: Boxplot of the ESTIMATE immune infiltration scores of samples across all the pre- and on-treatment datasets. The significances of the comparisons were from the Wilcoxon rank sum test.

**Fig. S7**: Response prediction was not improved by integrating our approach and TIDE.

**Fig. S8**: The hierarchical clustering of cancer types using the all the 2019 overlapping genes across the 31 TCGA cancer types, revealing that cancer types in related tissue lineages were clustered together.

## Supplementary Table Legends

**Table S1**: MHC I network association prediction of all the genes in the gene network.

**Table S2**: The selected positive and negative signature genes of the anti-PD-1 response for SKCM.

**Table S3**: Top 10% genes in our MHC I-association prediction, whose expression levels were significantly correlated with the immune infiltration score (absolute correlation ≥ 0.2) in the ???31 TCGA cancer types.

**Table S4**: Pathway enrichment analysis of the overlapping genes (Top 10% genes in our MHC I-association prediction, whose expression levels were significantly correlated with the immune infiltration score) in in the 34 TCGA cancer types.

**Table S5**: Target genes of 36 compounds that have been in clinical trials and already used for immuno (anti-PD-1)-targeted combination therapies in cancer treatment. The 36 compounds were collected from several review papers (see the main text). The known target genes of these compounds were compiled from the DGIdb database.

**Table S6**: Patient information of the 354 pre- or on-treatment tumor samples (anti-PD-1 or combination of anti-PD-1 and anti-CTLA-4) used for evaluating the prediction performance of the MIAS score.

