## Supplemental figures for "A computational network approach to identify predictive biomarkers and therapeutic combinations for anti-PD-1 immunotherapy in cancer"

### Slide 1
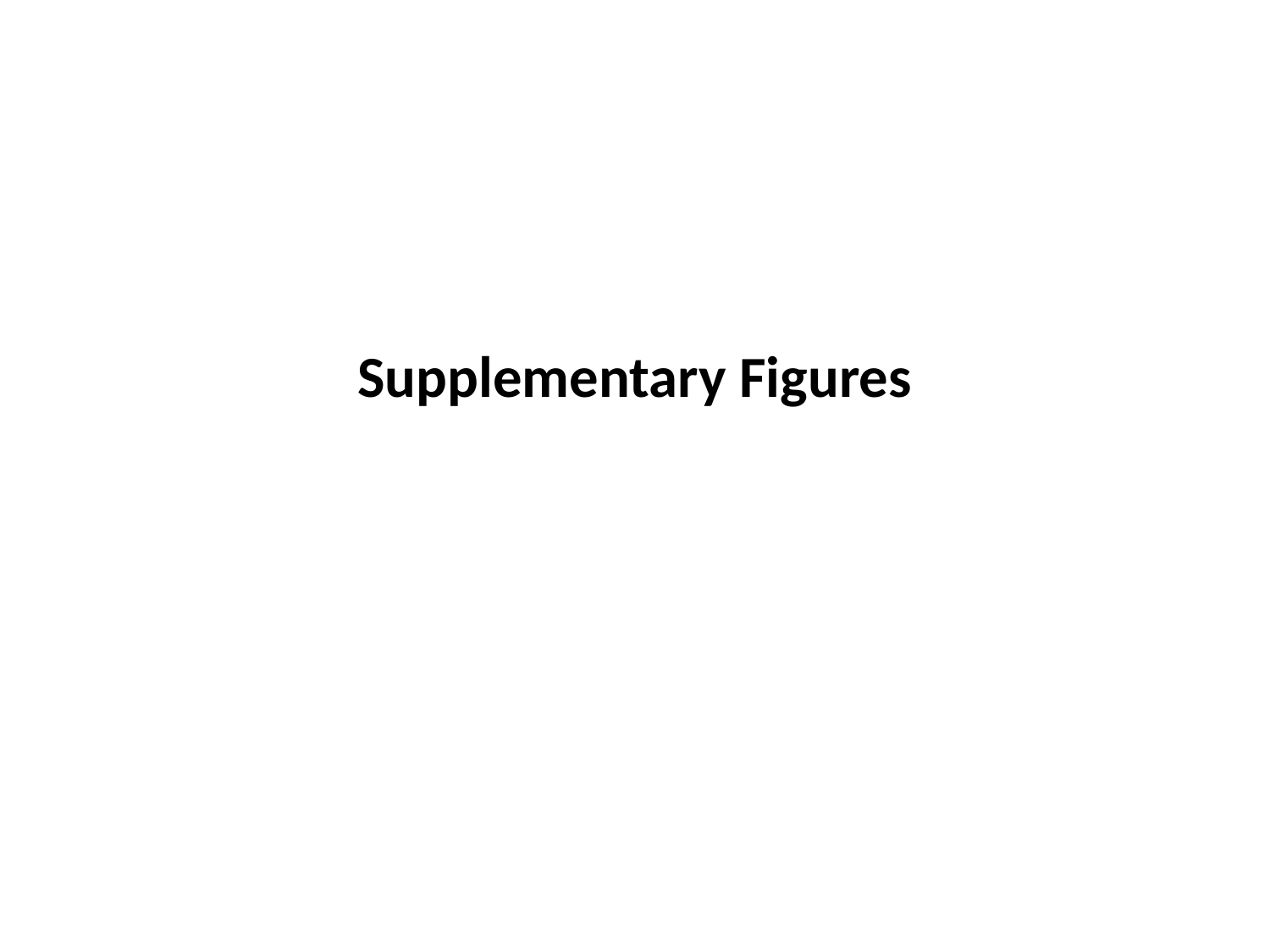

Supplementary Figures

### Slide 2
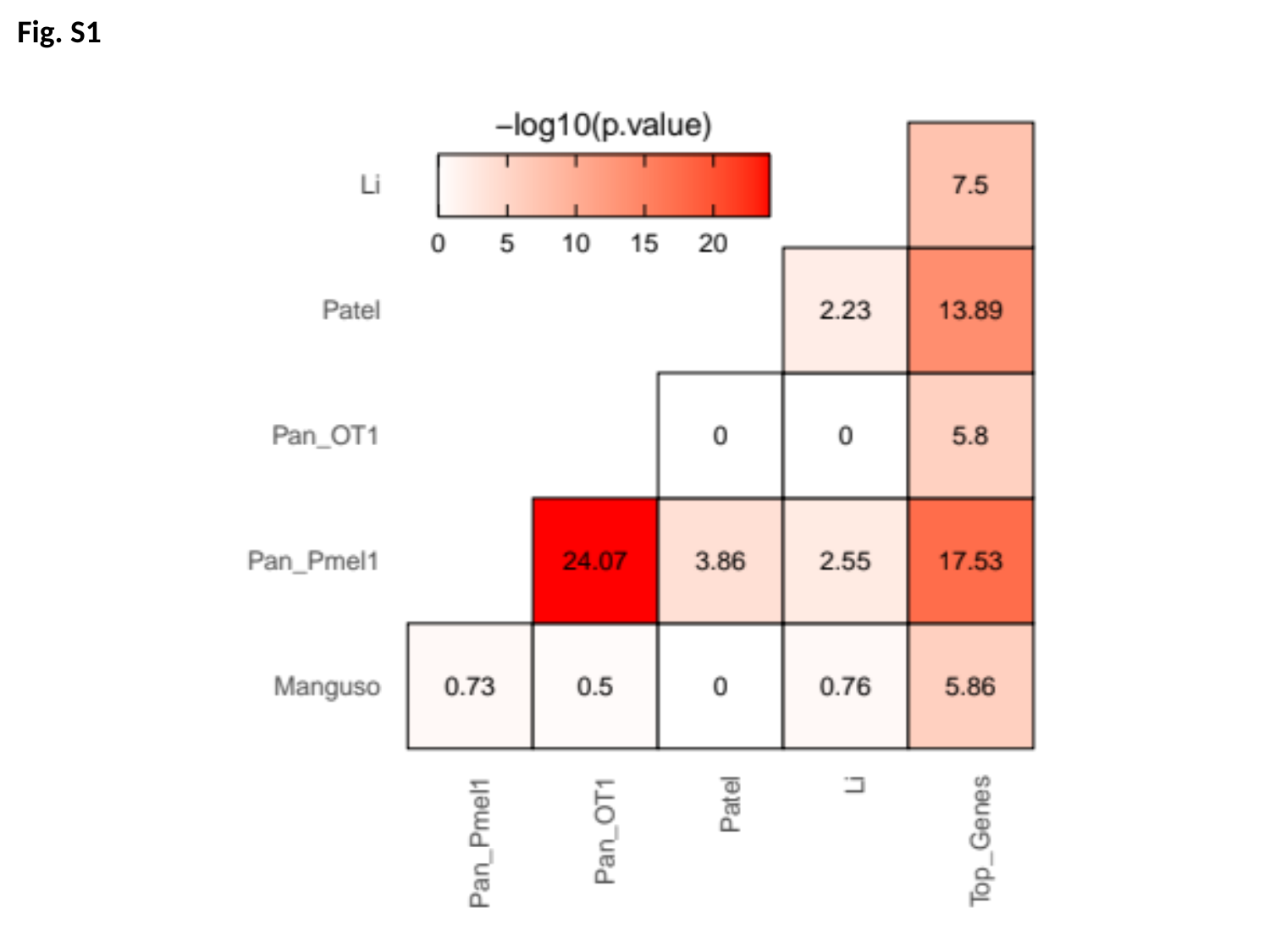

Fig. S1

### Slide 3
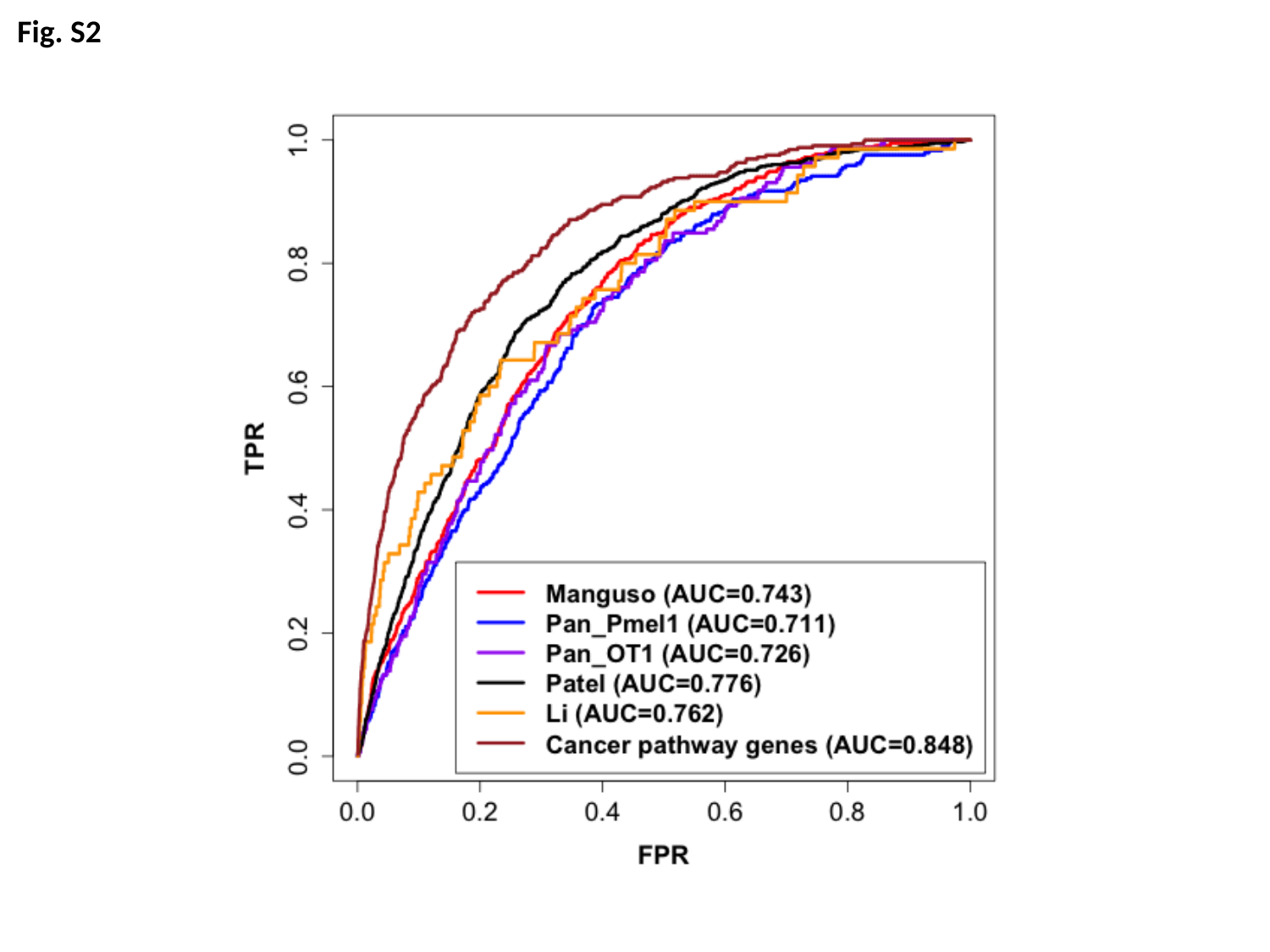

Fig. S2

### Slide 4
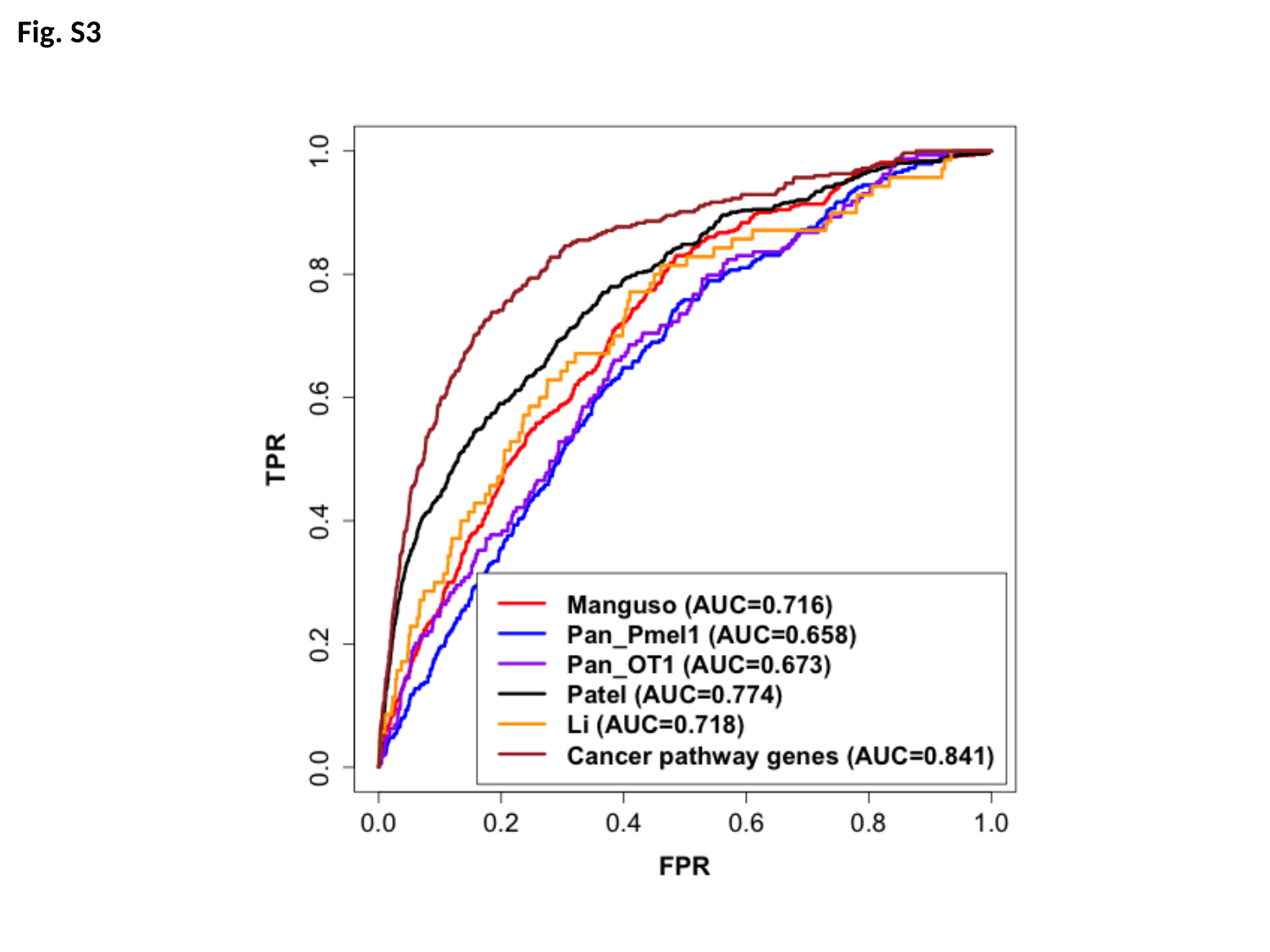

Fig. S3

### Slide 5
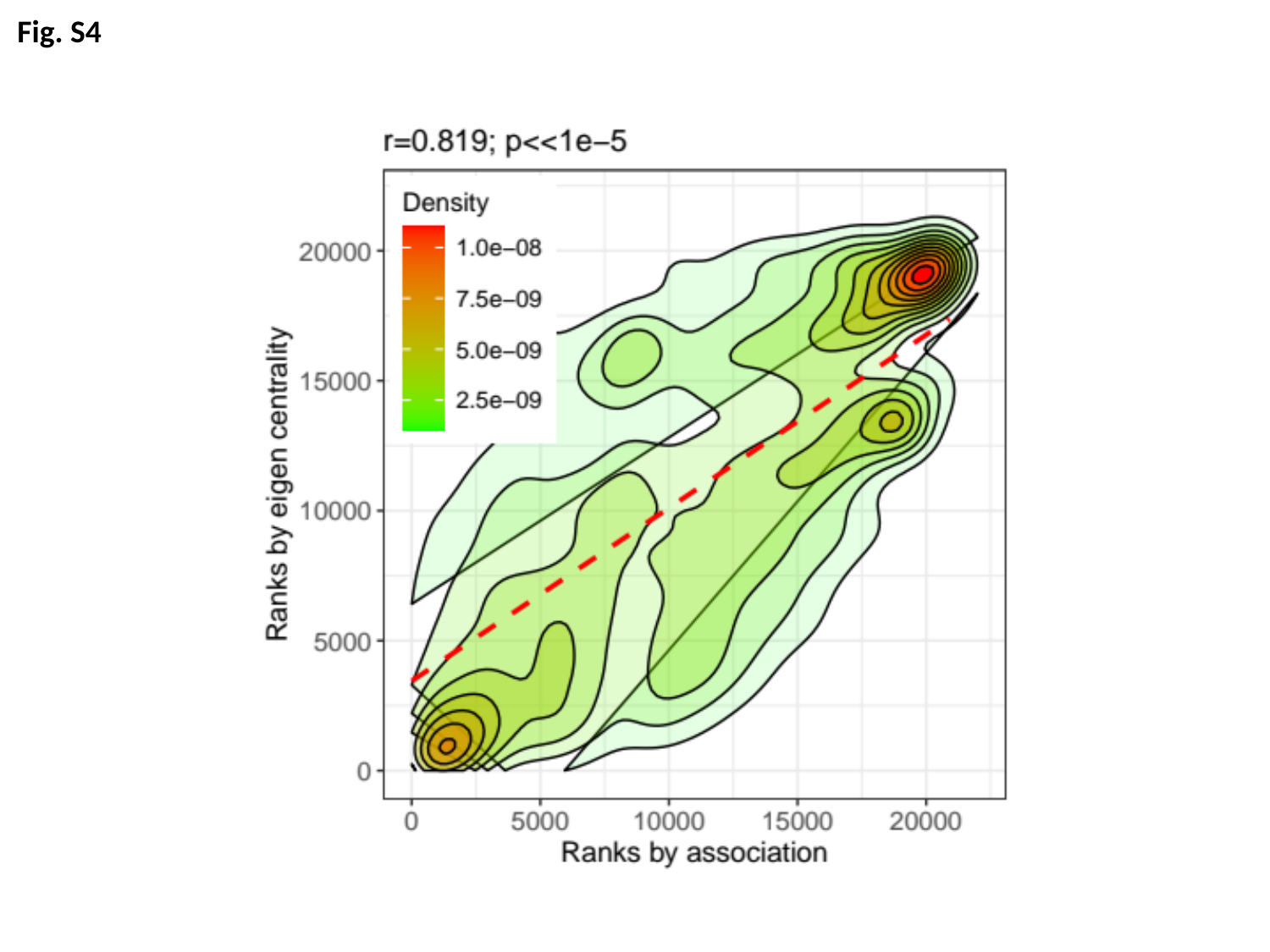

Fig. S4

### Slide 6
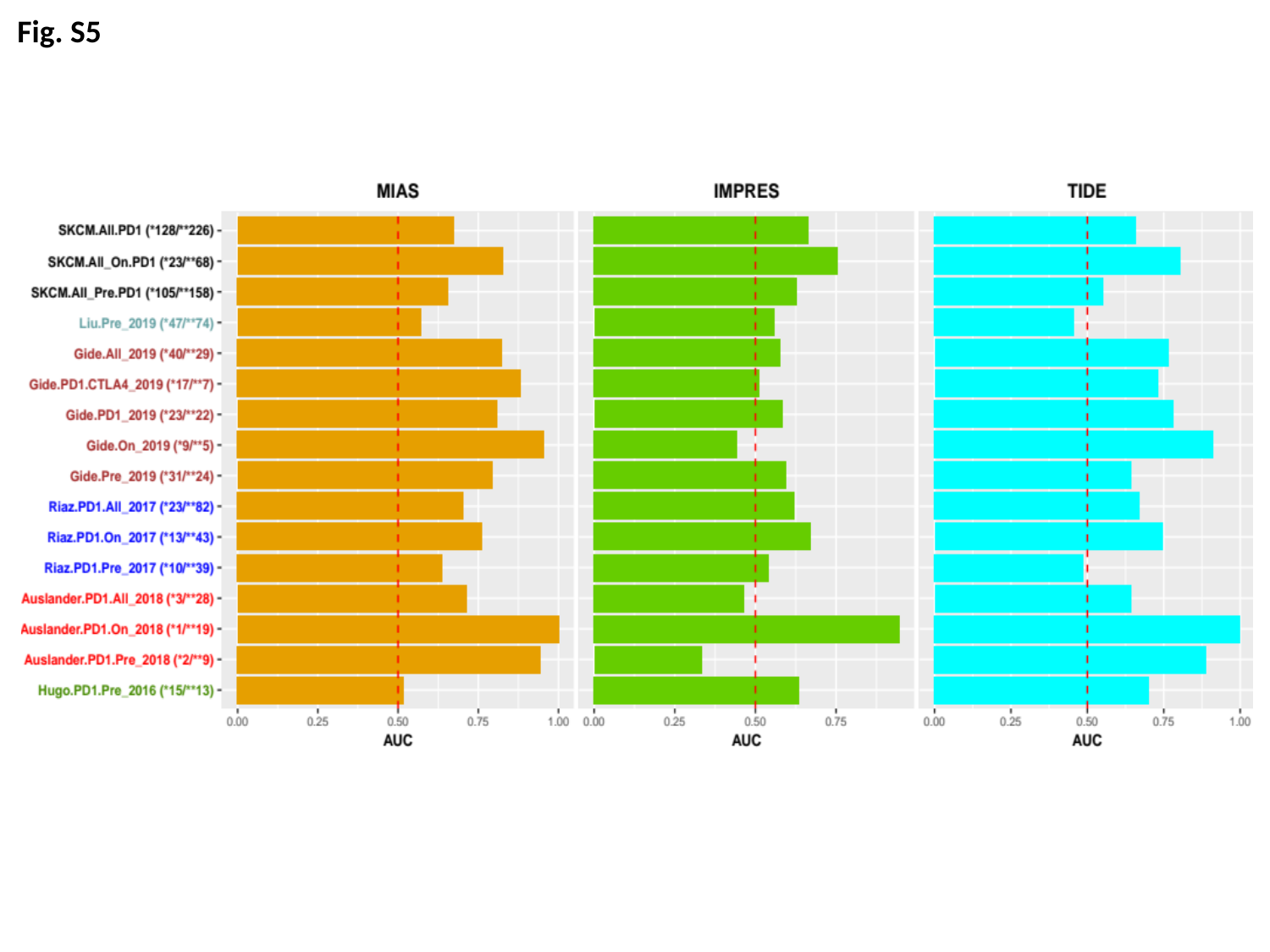

Fig. S5

### Slide 7
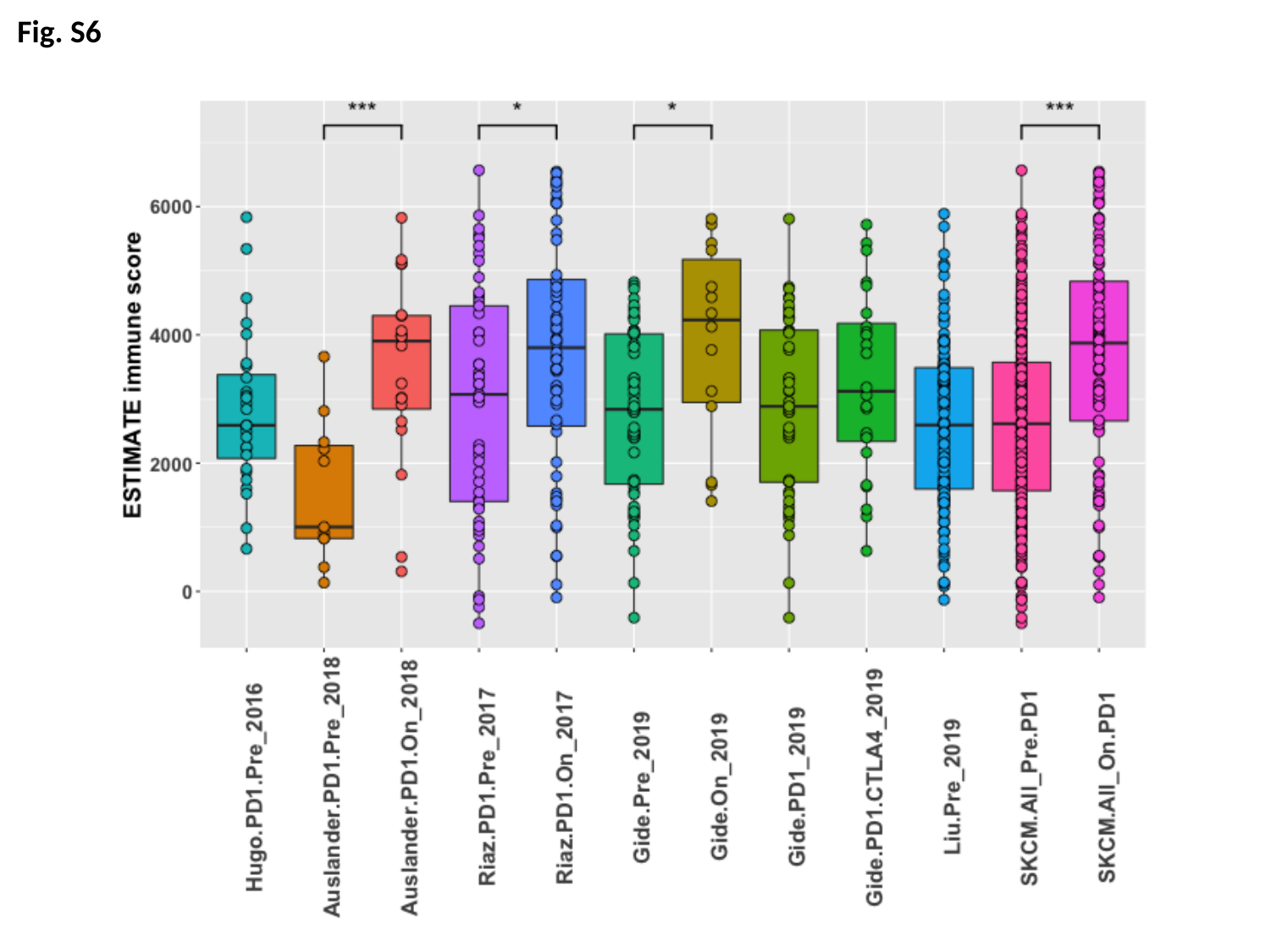

Fig. S6

### Slide 8
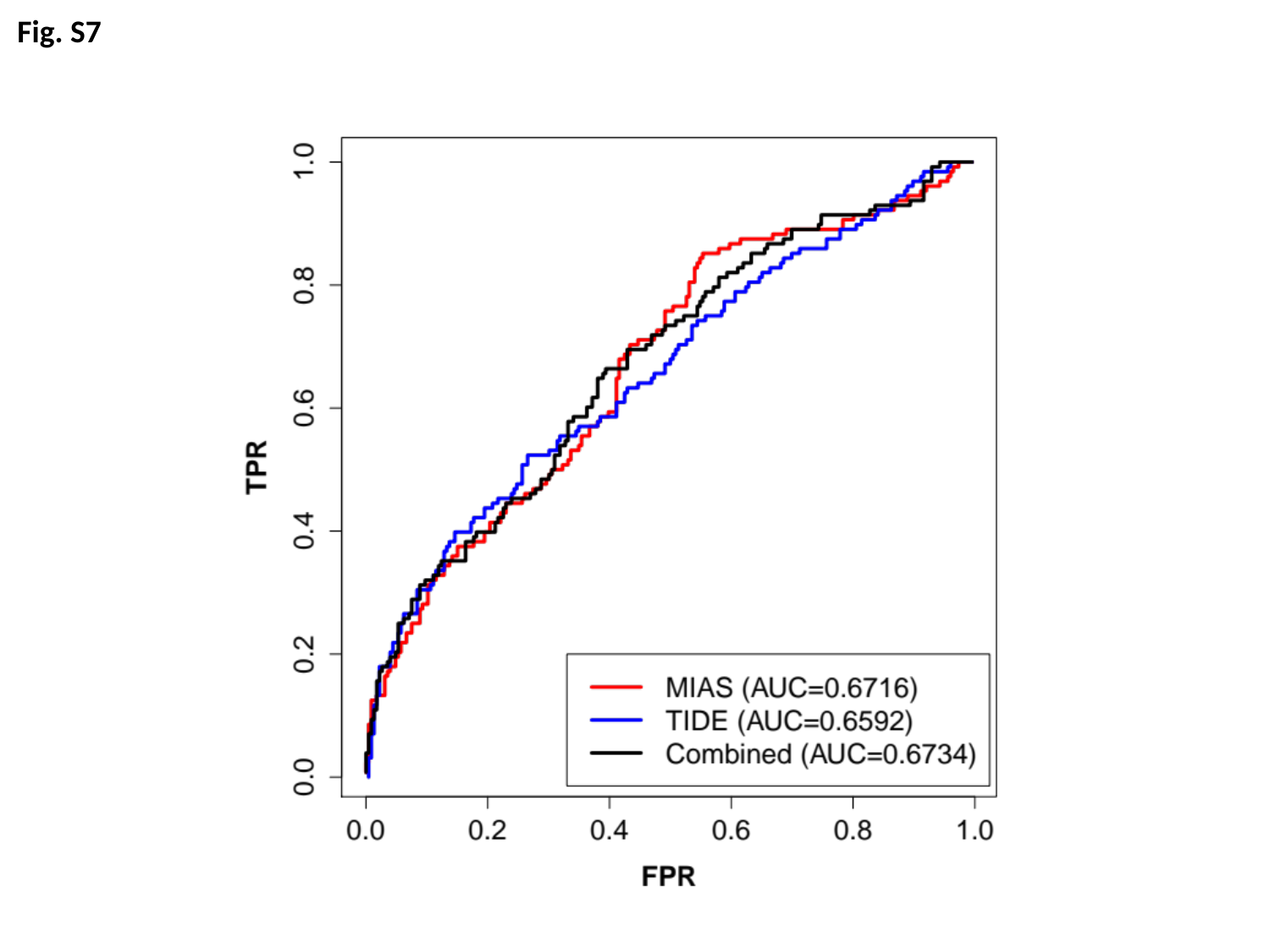

Fig. S7

### Slide 9
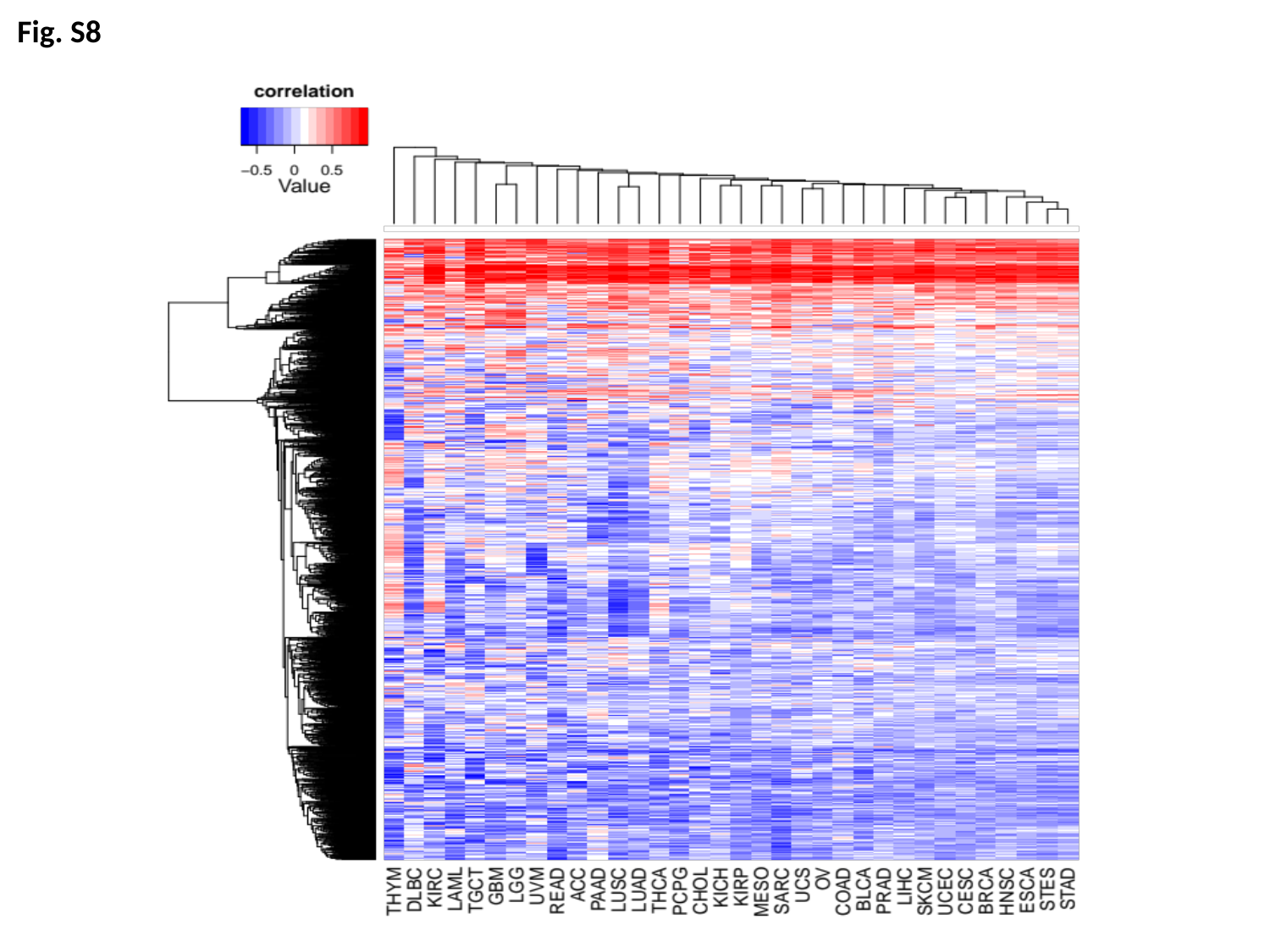

Fig. S8
